# Water exchange rates measure active transport and homeostasis in neural tissue

**DOI:** 10.1101/2022.09.23.483116

**Authors:** Nathan H. Williamson, Rea Ravin, Teddy X. Cai, Melanie Falgairolle, Michael J. O’Donovan, Peter J. Basser

**Affiliations:** Eunice Kennedy Shriver National Institute of Child Health and Human Development, National Institutes of Health, Bethesda, MD, USA; National Institute of General Medical Sciences, National Institutes of Health, Bethesda, Maryland, USA; Celoptics, Rockville, MD, USA; Wellcome Centre for Integrative Neuroimaging, FMRIB, Nuffield Department of Clinical Neurosciences, University of Oxford, Oxford, UK; National Institute of Neurological Disorders and Stroke, National Institutes of Health, Bethesda, MD, USA; National Center for Complementary and Integrative Health, National Institutes of Health, Bethesda, MD, USA

**Author notes:** N.H.W. and R.R. contributed equally to this work.

**Keywords:** Water co-transport, Stroke, Tissue microstructure, Diffusion exchange spectroscopy (DEXSY), Porous media, Membrane permeability, Transcytolemmal water exchange, NMR-MOUSE, Biomarkers

## Abstract

For its size, the brain is the most metabolically active organ in the body. Most of its energy demand is used to maintain stable homeostatic physiological conditions. Altered homeostasis and active states are hallmarks of many diseases and disorders. Yet there is currently no reliable method to assess homeostasis and absolute basal activity or activity-dependent changes non-invasively. We propose a novel, high temporal resolution low-field, high-gradient diffusion exchange NMR method capable of directly measuring cellular metabolic activity via the rate constant for water exchange across cell membranes. Using viable *ex vivo* neonatal mouse spinal cords, we measure a component of the water exchange rate which is active, i.e., coupled to metabolic activity. We show that this water exchange rate is sensitive primarily to tissue homeostasis and viability and provides distinct functional information in contrast to the Apparent Diffusion Coefficient (ADC), which is sensitive primarily to tissue microstructure but not activity.

**SIGNIFICANCE STATEMENT:** Despite what physiology textbooks may report, water transport across membranes is not only a passive process. However, current understanding is limited because standard techniques can only measure net flux (the difference between water moving in and water moving out). Even so, water is constantly exchanging between the inside and outside of cells and organelles without net flux during homeostasis. We developed a Magnetic Resonance method able to “see” water molecules exchanging on shorter timescales than could be observed before. In neural tissue we find most water exchange is active, that is, linked to ATP-driven processes. This method may one day be translated to clinical MRI applications for measuring cellular function and activity in the human brain and body.

## Introduction

Current non-invasive methods used to detect activity in the central nervous system (CNS), such as BOLD fMRI, are indirect and sensitive only to metabolic *changes* (1), which account for just 5% of the total energy budget. The remaining 95% is devoted to homeostasis, i.e., maintaining steady-state activity levels conducive to normal brain function (2). A large field of resting-state fMRI is devoted to characterizing different brain states, but can only indirectly infer them based on relative BOLD signal changes (3).

Steady state means that intra- and extra-cellular concentrations and volumes remain relatively constant over time and that there is no significant net flow or flux of water and other species across cell and organelle membranes. While the steady state seems static, molecules, including water are constantly, actively exchanging between inside and outside the cell (4). Homeostasis can result from different water exchange rates as long as the transport in and out are the same. However, steady state water exchange is normally invisible to biophysical measurements. Here we show Nuclear Magnetic Resonance (NMR, or simply, MR) can readily follow steady state water exchange rapidly, and over time.

Previously accepted knowledge about water transport across biological membranes (5) has come under scrutiny in the light of new evidence (6). Previously, water was thought to transport only passively, e.g., by diffusing across lipid membranes, and through aquaporin water channels (7, 8). However, many studies report that membrane transport proteins actively cotransport hundreds of H_2_O molecules per ion or metabolite (9–14). With respect to the CNS, active water transport is known to be important for the function of epithelial tissue, namely in the choroid plexus, for which water flux measurements suffice (6). Active water transport within parenchymal nervous tissue has not been characterized, in part due to the dearth of techniques capable of measuring it under steady-state conditions. In fact, it may only be possible to do noninvasively with NMR.

NMR can measure the rate constant for steady-state tran-scytolemmal water exchange, hereafter called the “exchange rate”. NMR does so by directly encoding and measuring signals from protons (^1^H) on water, and following their evolution as intra- and extra-cellular water pools mix. NMR findings suggest that there is a component of water exchange linked to Na^+^*/*K^+^–ATPase activity (15–21). Continual Na^+^*/*K^+^–ATPase activity accounts for about half the energy budget of CNS tissue (22) — approximately 2 ×10^5^ ATP molecules per second per *µ*m^3^ of neural tissue (22). If each cycle of a Na^+^*/*K^+^ pump cotransports *>*100 H_2_O molecules, then more than 2×10^7^ water molecules exchange per second per *µ*m^3^. The steady-state water exchange rate amplifies the steady-state ion transport rate. Then, the exchange rate can potentially be used as a marker to assess cellular energetic and homeostatic states.

Traditionally, intracellular and extracellular water pools have been distinguished based on their differing relaxation rates by introducing contrast agents to dope the water in the extracellular space using a method called Dynamic Contrast Enhanced (DCE) Magnetic Resonance Imaging (MRI) (23). DCE MRI has been widely used to measure active water exchange (4, 15–19). However, this method has the disadvantage of requiring exogenous contrast agents, usually containing gadolinium (Gd), which typically are excluded from the brain by the blood-brain barrier (BBB) (16), and are increasingly associated with some forms of toxicity (24), long-term accumulation in tissue (25), and even environmental contamination (26). Another MR method uses diffusion measurements to distinguish between intracellular and extracellular water based on their differing mobilities or apparent diffusivities and observes exchange by varying the timescale of diffusion encoding (27, 28). This method was recently shown to be sensitive to active water exchange (20, 21). However, effects of restriction and exchange both vary with diffusion encoding time and are difficult to separate using one-dimensional diffusion measurements (29).

A promising alternative MR method for measuring active water exchange is Diffusion Exchange SpectroscopY (DEXSY) (30). DEXSY is a multidimensional MR method (31) that combines two diffusion encoding times separated by a mixing time which can be varied independently to isolate and measure exchange unaffected by restriction (32, 33), overcoming limitations of one-dimensional diffusion measurements. DEXSY and variants of DEXSY have been used to non-invasively measure water exchange rates using only the endogenous tissue water as a reporter, without requiring the introduction of exogenous contrast agents, making this methodology amenable to translational MR imaging applications without the attendant disadvantages of DCE (32–42).

Here we study the activity dependence of the exchange rate measured by DEXSY. A low-field single-sided MR test system was previously developed for studying activity dependence of MR parameters (43). Viable “live” *ex vivo* CNS tissue models were used because their condition could be precisely controlled and perturbed and they are free of blood flow, respiratory, and motion artifacts. We have made significant improvements, having increased the SNR by an order of magnitude by using a specially built solenoid radiofrequency (RF) coil paired with *ex vivo* neonatal mouse spinal cords in order to provide a high filling factor (41). A full DEXSY acquisition required many scans and > 1 hr per exchange rate measurement (41, 44). We have reduced the time to as little as 1-10 min by while separating diffusion, relaxation, and exchange effects in the multidimensional DEXSY measurement (32, 42). We can now report exchange rates in realtime on a biological timescale on living specimen. Finally, we harness the strong static gradient (15.3 T/m) to perform diffusion and DEXSY measurements with sub-millisecond diffusion encoding times and sub-micron spatial resolution (41). In contrast, conventional diffusion MR measurements are performed at high-field using gradient coils to apply time-varying pulsed field gradients (PFG) with considerably lower gradient strengths and longer diffusion encoding times (45). “Low-field, high-gradient” DEXSY allows us to measure exchange processes which are too fast to be observed with DCE or conventional PFG methods.

N.B. MRI exchange measurements may not be necessary to quantify steady-state baseline levels of CNS homeostasis. An NMR measurement averaged over a large region of interest (ROI) in the brain parenchyma may be sufficient to observe physiological and pathological steady-state conditions. As a first step, we perform DEXSY NMR with the thought that it can be combined with MRI in the future, as was done for other diffusion and spectroscopic NMR sequences (46). This study also provides a new direction for studying water homeostasis in the CNS, in particular revealing steady-state water cotransport with low-field, high-gradient DEXSY NMR.

In order to find the active component of exchange and its relevance to homeostasis we use the following three experimental perturbations. We first use the effect of temperature on live vs. fixed tissue. In live tissue, the exchange rate is thought to be a sum of active and passive processes occurring in parallel, *k* = *k*_*a*_ + *k*_*p*_ (19). Fixed tissue, is by definition at equilibrium, implying no active exchange, and therefore the steady-state exchange is entirely passive. We find that *k* has a stronger dependence on temperature in live tissue than in fixed tissue, indicating that *k*_*a*_ exists and is measurable with low-field, high-gradient DEXSY. Second, we inhibit Na^+^*/*K^+^ –ATPase activity with ouabain. This decreases the water exchange rate by 70%, demonstrating that active water exchange is linked to ion transport. Lastly, we study the effects of oxygen and glucose deprivation as a model for ischemia/stroke. Results collectively suggest that the exchange rate is a measure of tissue viability. This technique can be used for detecting the onset of pathological states. From concurrent diffusion measurements, we find that the ADC has a similar dependence on temperature for live and fixed tissue, indicating that diffusion measurements are not directly sensitive to metabolic activity. Instead, we find that the ADC follows structural changes not necessarily correlated to the exchange rate.

## Results

### Exchange rates are linked to the non-equilibrium metabolic activity in live tissue

Exchange rates were measured in real-time on *n* = 6 fixed and *n* = 7 live *ex vivo* neonatal mouse spinal cords undergoing step changes in temperature: 25 → 7 → 25 → 35 → 25°C (Fig. 1a and b). Four of the seven live samples only underwent the first 25 → 7 → 25°C portion of the temperature variation protocol because holding the system at 35°C for 40 minutes of measurements was challenging for the spinal cord specimen and caused the exchange rate to run down, indicative of reduced tissue viability. Temperature changes had a greater effect on the exchange rate for live samples than for fixed ones. At 7°C, exchange rates were similar between live and fixed samples whereas at 25°C and 35°C, exchange rates were greater for live than for fixed samples.

The 25°C condition was repeated in a “test/re-test” manner to check that samples recovered from being subjected to 7°C and 35°C. Exchange rates for fixed samples consistently returned to the same values during the 2^nd^ and 3^rd^ 25°C. In contrast, live samples showed decreased exchange rates at the 3^rd^ 25°C due to rundown during the 35°C condition.

**Fig. 1.**
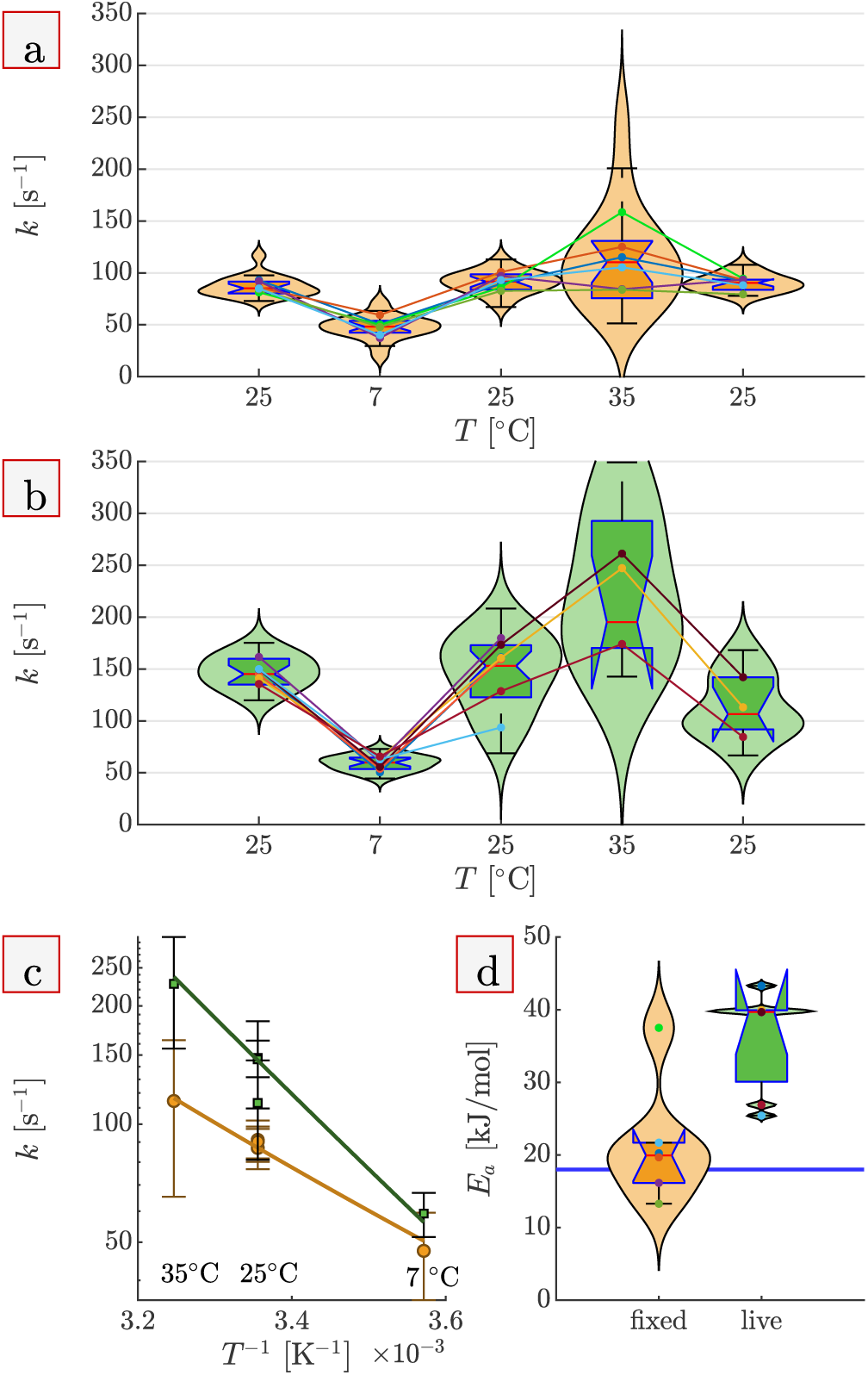
Temperature dependence reveals active, steady-state water exchange. Exchange rates, *k*, for fixed (a) and live (b) spinal cords measured at different temperatures. Box and violin plots include all *k* measurements (three measurements per sample) and are discussed in the Methods. Connected dots show the mean *k* at each temperature and are colored differently for each sample. (c) Arrhenius plot of the dependence of *k* on *T* ^−1^ (the inverse of the absolute temperature) for fixed (orange) and live (green) spinal cords. Mean values (symbols) and standard deviations (whiskers) for all fixed and live samples at each temperature condition are presented. Data points for the 1^st^, 2^nd^, and 3^rd^ 25° C conditions are shown separately. Lines show the mean of Arrhenius model fits. Slopes of the lines are proportional to *E*_*a*_. (d) Boxplots comparing activation energies (*E*_*a*_) between fixed and live spinal cords. Dots show the values of *E*_*a*_ for each sample and use the same colors as in (a) and (b). *E*_*a*_ = 21 *±* 8 kJ*/*mol (mean ± SD) for fixed samples, similar to *E*_*a*_ = 18 found for water self-diffusion in artificial cerebrospinal fluid (aCSF, solid blue line, see also Fig. 2). *E*_*a*_ = 36 *±*7 kJ*/*mol for live spinal cords and is significantly greater than *E*_*a*_ for fixed spinal cords (*p* = 0.005). Therefore, active exchange exists.

An Arrhenius plot of the exchange rates vs. the inverse absolute temperature is shown in Fig. 1c. The slope of the logarithm of *k* vs. *T* ^*−*1^ is proportional to the activation energy (*E*_*a*_). The slope is steeper (i.e., *E*_*a*_ is greater) for live spinal cords than for fixed spinal cords. *E*_*a*_ values for each sample were estimated by fitting the data with an Arrhenius model, *k* = *A* exp(− *E*_*a*_*/RT*) where *A* is a prefactor, *R* is the ideal gas constant, and *T* is the absolute temperature. *E*_*a*_ estimates are compared between fixed and live samples in Fig. 1d. For fixed spinal cords, the *E*_*a*_ is similar to the *E*_*a*_ for self-diffusion of pure water (47). This affirms that exchange in fixed tissue is driven by equilibrium thermal energy and mediated by passive water permeability through openings, such as pores or channels in the membrane (48). In comparison, *E*_*a*_ for live spinal cords is significantly greater than the value for fixed ones. Exchange rate rundown at 35°C leads to *E*_*a*_ of live tissue being underestimated. Therefore, water exchange in live tissue is not only driven by thermal energy — it must also be driven by non-equilibrium active processes.

### Apparent diffusion coefficients are sensitive to tissue microstructure but not activity

Water diffusion in the *y*-direction, perpendicular to the orientation of the spinal cord, was also measured in real-time on the *n* = 6 fixed and *n* = 7 live spinal cords at 25°, 7°, 25°, 35°, and 25°C. Raw signals are shown in Fig. S2. Apparent Diffusion Coefficients (ADC_*y*_) were estimated from the initial signal decay (*b* = 0.096 to 1.5 ms*/µ*m^2^). ADC_*y*_ consistently recovered upon returning to 25°C (Fig. 2a and b), including for live spinal cords at the 3^rd^ 25°C which showed decreased exchange rates due to rundown after the 35°C condition. ADC_*y*_ values for fixed spinal cords are significantly less than for live spinal cords (Fig. 2c).

**Fig. 2.**
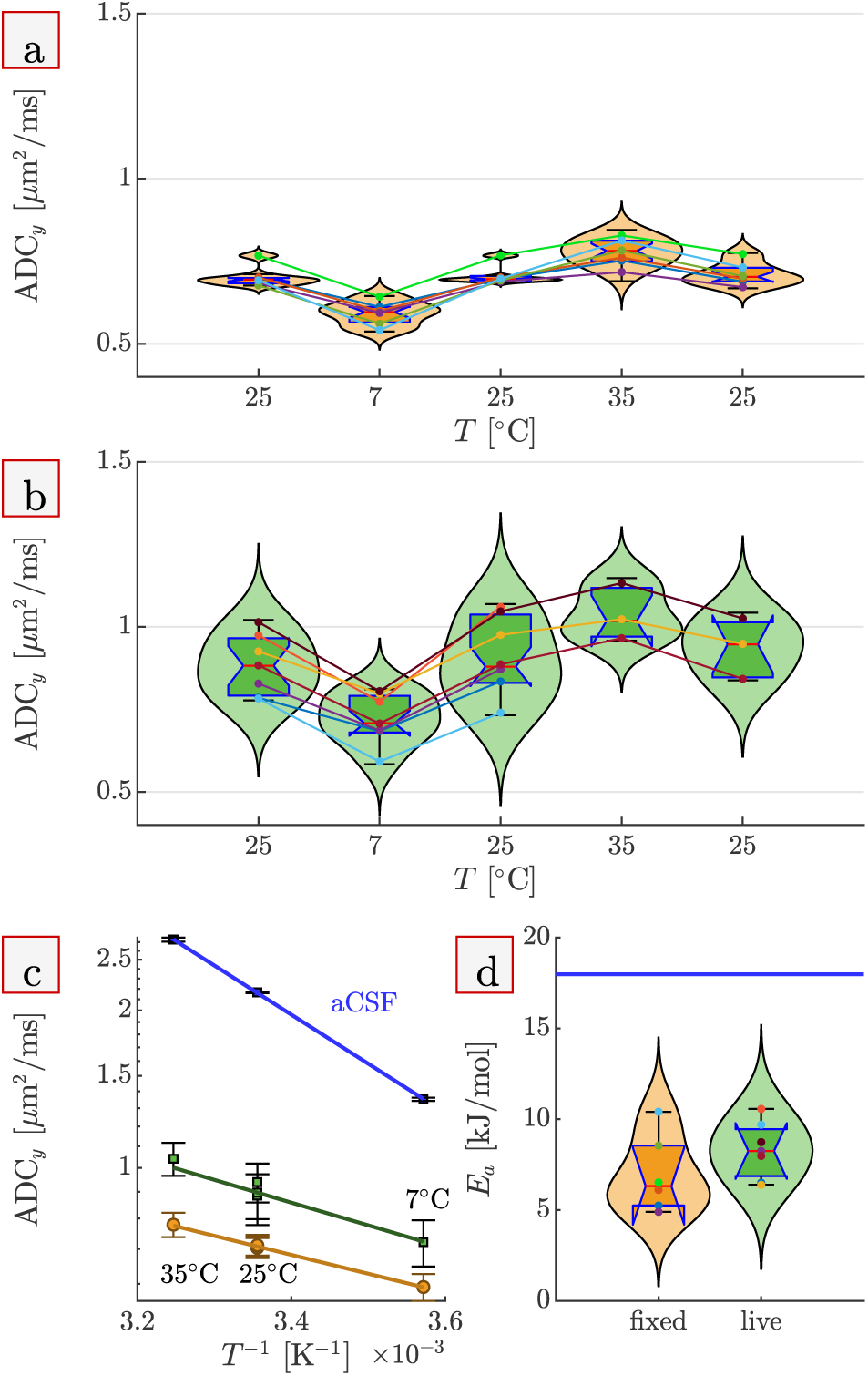
Temperature dependence of ADC_*y*_. Apparent diffusion coefficients of fixed (a) and live (b) spinal cords at the different temperatures. c) Arrhenius plot of ADC_*y*_ at each temperature condition for fixed (orange) and live (green) spinal cords, as well as free diffusion coefficients of pure aCSF (blue). d) Activation energies (*E*_*a*_) of water diffusion for fixed and live spinal cords (boxplots) and aCSF (solid blue line). Dots show *E*_*a*_ values for each sample using the same colors as in (a) and (b). A greater description of plot details is provided in the caption of Fig. 1. For aCSF, *E*_*a*_ = 18 kJ*/*mol, similar to values reported for pure water over the same temperature range *E*_*a*_ = 18 *−* 20 kJ*/*mol (47). *E*_*a*_ of ADC_*y*_ is not significantly different between live (*E*_*a*_ = 8.3 *±* 1.5 kJ*/*mol), and fixed spinal cords (*E*_*a*_ = 7.0 *±* 2.1 kJ*/*mol), (*p* = 0.21).

Diffusion coefficients of pure artificial cerebrospinal fluid (aCSF show an Arrhenius temperature dependence consistent with literature reports for pure water (47) (Fig. 2c and d). Water in aCSF diffuses freely (unimpeded by restriction of membranes) such that the thermal energy translates directly to increased mean-squared displacements. ADC_*y*_ of fixed and live spinal cords were less affected by temperature. This is because water experiences more interactions with membranes as the temperature increases, retarding it and keeping the ADC_y_ from increasing as much as it would if it were free. This affirms that ADC_*y*_ probes hindrances and restrictions imposed by tissue microstructure. *E*_*a*_ of ADC_y_ for live and fixed spinal cords are not significantly different. Therefore, the measured ADC is insensitive to non-equilibrium metabolic activity.

In addition to exchange rates, DEXSY experiments also provide diffusion-weighted spin-lattice relaxation rates (*R*_1 DW_). *R*_1_ is the reciprocal of *T*_1_ and is inversely related to its rotational mobility (49). Higher temperature increases rotational mobility and decreases *R*_1_. The strong diffusion weighting (*b* = 4.5 *ms/µm*^2^) filters out signal based on translational mobility such that *R*_1 DW_ is associated with water which is more hindered and restricted by membranes within the tissue. *R*_1 DW_ for fixed spinal cords is higher than for live spinal cords (Fig. S3). *R*_1 DW_ recovered for fixed and live spinal cords when returned to 25°C after 7°C and 35°C. *E*_*a*_ of 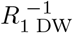 for live spinal cords are greater than for fixed spinal cords when based on a t-test (p<0.05) but are not significantly different when based on the 95% confidence interval (CI) of median values (Fig. S3c and d).

### Active water exchange is linked to ion transport

While a comparison of *E*_*a*_ values between live and fixed tissue indicates that water exchange is linked to active cellular processes, it does not reveal a link to specific enzymes or specific cellular processes. We tested the role of Na^+^*/*K^+^–ATPase by measuring water exchange in real-time during the addition of 100 *µ*M ouabain on *n* = 3 spinal cords (Fig. 3). 100 *µ*M ouabain is expected to reduce Na^+^*/*K^+^–ATPase activity by 65% and the effect increases by only 6% at higher doses (50, 51). 100 *µ*M ouabain took effect roughly 10 minutes after treatment. The exchange rate dropped by 71*±* 8%, indicating that the majority of water exchange is linked to ion transport. The apparent diffusion coefficient decreased by 9*±* 1%, and restricted water fraction (*f*) increased by 22 *±* 6% (see Fig. S4), consistent with cells swelling due to a net influx of ions and water (52).

**Fig. 3.**
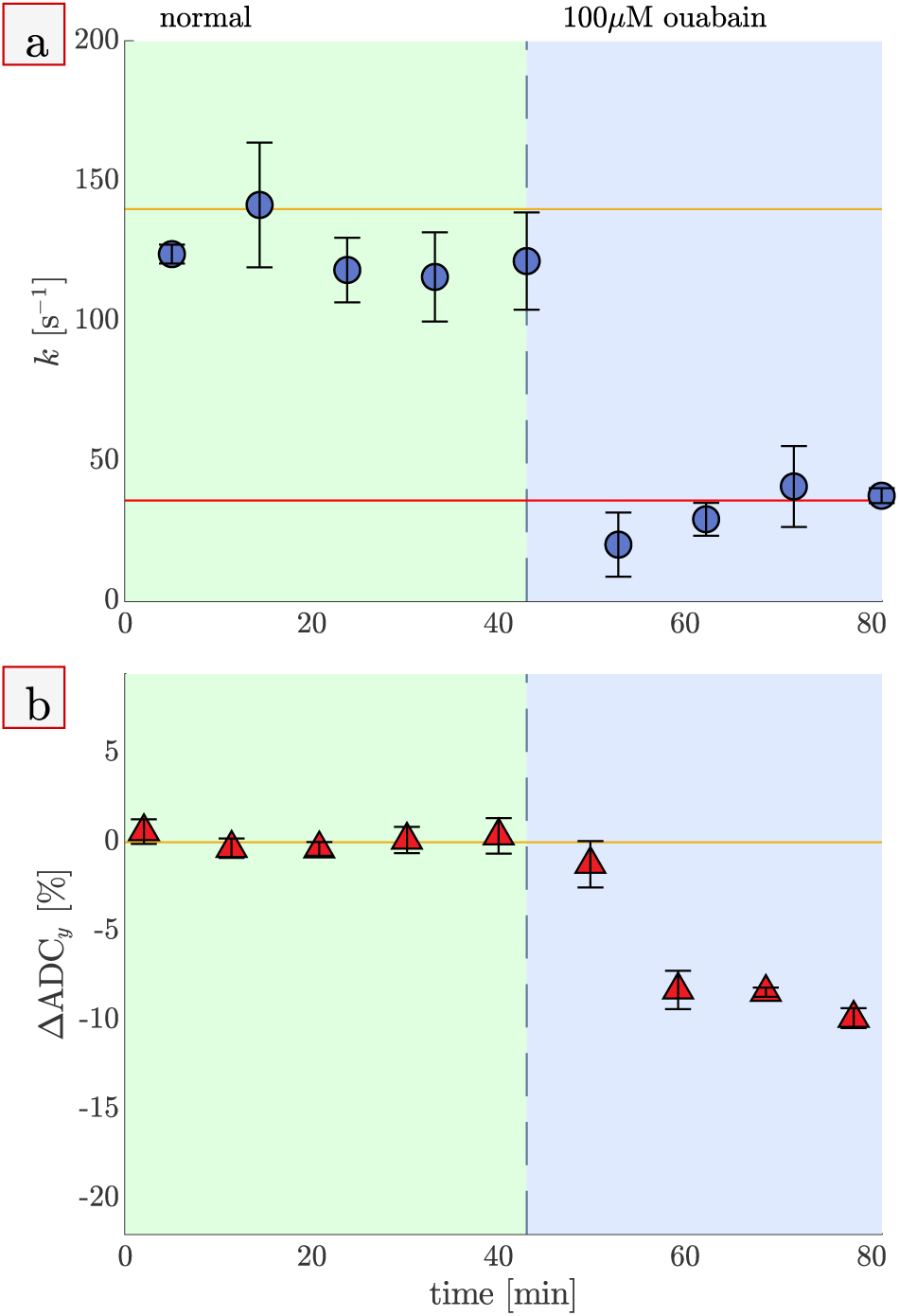
Exchange rates drop by 71% after inhibiting the sodium–potassium pump with ouabain. Real-time exchange rate (a) and percentage change in apparent diffusion coefficient from baseline (b) measurements on live spinal cords under normal condition (green shading) and after (magenta) addition of 100 *µ*M ouabain at 25°C. Mean values (circles) and standard deviations (whiskers) from n=3 samples are presented. In (a), the solid orange line shows the average exchange rate value from all normal conditions in this paper (*k* = 140 s^*−*1^). The red line shows the mean value after ouabain addition (*k* = 36 s^−1^). (See Fig. S4 for associated *f* and *R*_1 DW_ data).

### Stroke model suggests water exchange rates measure tissue viability

Cellular damage during stroke is linked to the duration of hypoxia or hypoglycemia. We investigated the potential that the exchange rate could be used to monitor reduced tissue viability as a result of stroke by comparing the effect of a shorter 40 min (*n* = 8, Fig. 4a, c, and e) and a longer 70 min (*n* = 9, Fig. 4b, d, and f) oxygen–glucose deprivation (OGD) model.

For all samples, pO_2_ reduced quickly after the switch and recovered when washing back, confirmed by monitoring *R*_1_ (Fig. 4a and b). ADC_*y*_ first dropped within 10 minutes of the switch to OGD and dropped a second time after roughly 30–40 minutes (Fig. 4c and d). After switching back to normal aCSF, ADC_*y*_ increased slightly within 10 minutes and slowly recovered to baseline over the next two to three hours. In some cases, the ADC_*y*_ increased above baseline (see examples in Fig. S9). The exchange rate behaved differently from ADC_*y*_. After switching to OGD, the exchange rate first dropped after roughly 30–40 minutes (Fig. 4e and f). In the 40 min protocol, the samples were switched back to normal aCSF while the exchange rate was still dropping, rescuing some (but not all) of the samples and preventing their exchange rate from decreasing further. In the 70 minute protocol, the switch back to normal aCSF came after the exchange rate fully dropped. Exchange rates at the end of the 70 minute OGD condition were similar to exchange rates measured on ouabain-treated samples. The standard deviations are smallest at this point, perhaps because all samples were inactive, i.e., had reached complete energetic failure. After returning back to normal aCSF, samples recovered only slightly, if at all, and never back to baseline.

**Fig. 4.**
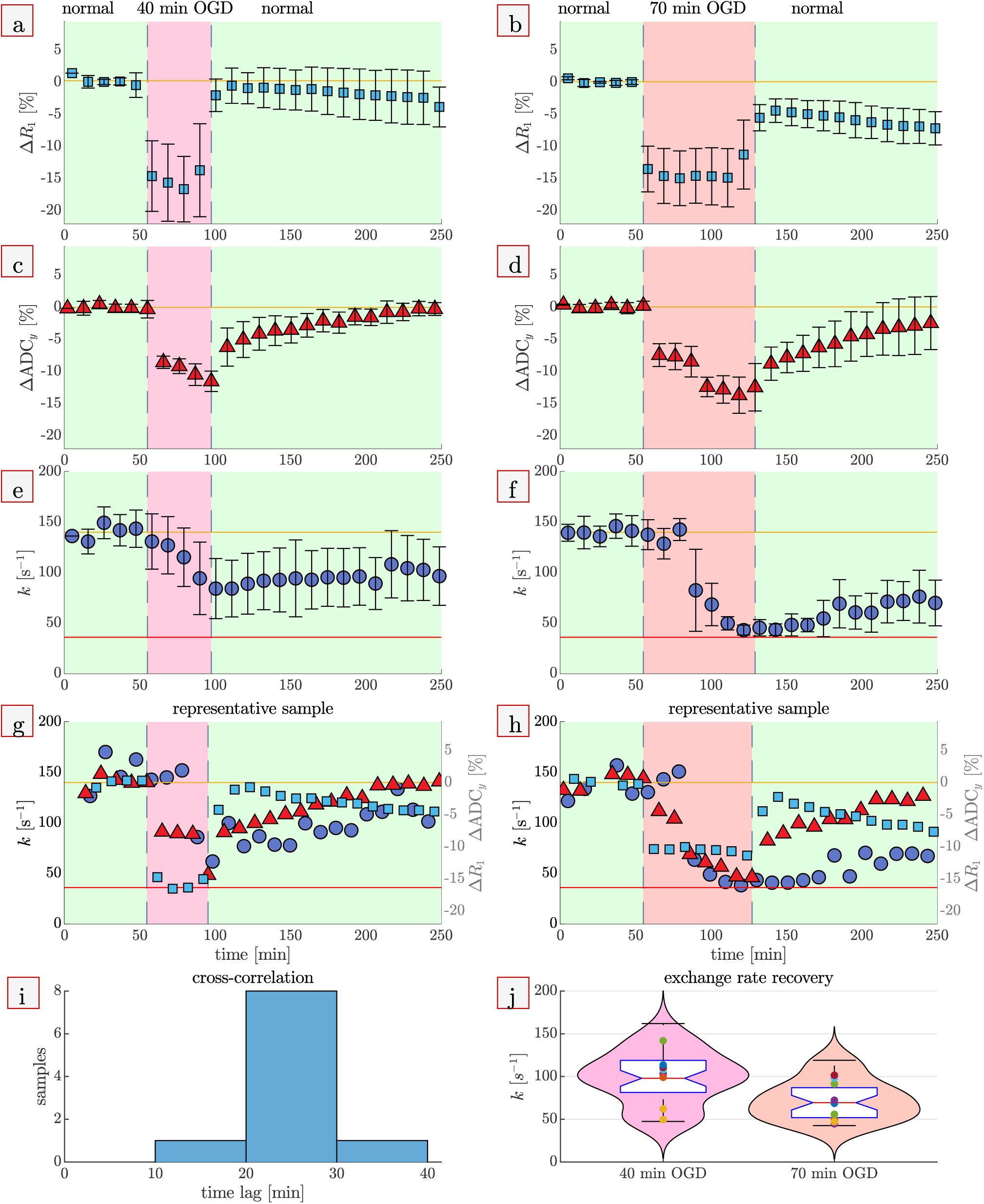
Exchange rates are highly regulated until viability is compromised during oxygen and glucose deprivation. Recordings of *R*_1_, ADC_*y*_, and *k* while switching from normal media bubbled with 95% pO_2_ to glucose-free media bubbled with 1% pO_2_ and washing back to normal after 40 minutes (left column) or 70 minutes (right column), at 25°C. a–f) Time series averages show means (symbols) and standard deviations (whiskers) of n=8 samples for 40 minute OGD and of n=9 for 70 minute OGD. g, h) Time series for representative samples, with *R*_1_ (light blue squares) and ADC_*y*_ (red triangles) values associated with the right-hand y-axis and exchange rate (dark blue circles) values associated with the left-hand y-axis. i) Histogram of how long the exchange rate response lags behind ADC_*y*_, calculated from cross-correlation analysis on the ten samples for which ADC_*y*_ and exchange rate are significantly correlated. j) Box and violin plots comparing the exchange rate values during recovery from 40 min OGD and 70 min OGD (averaged over the period 40–70 min after switching back to normal aCSF). Time series and Pearson correlations for all samples are shown in Fig. S9.

For a passive system it is expected that the ADC and exchange rate will be correlated through changes in cell volume (See Fig. S6). However, in light of active exchange, the ADC and the exchange rate may not necessarily be correlated and may be linked to system characteristics which are independently regulated. We find that the ADC_*y*_ and the exchange rate are significantly correlated for only 10 out of 17 samples (*p <* 0.05, Fig. S9). The correlation originates from the second drop of ADC_*y*_ aligning with the only drop of exchange rate. The first drop of ADC_*y*_ occurs while exchange rate is unaffected. A cross-correlation analysis confirmed that the exchange rate was first affected 20 to 30 minutes after the ADC_*y*_ first dropped (Fig. 4i). When washing back to normal aCSF, ADC_*y*_ recovered to (or overshot) baseline whereas exchange rates did not. Therefore, ADC and exchange rate are not necessarily correlated and must provide independent information. Exchange rates averaged over the period 40–70 min after switching back to normal aCSF were significantly higher for the 40 min protocol than for the 70 minute protocol (Fig. 4j), suggesting that the exchange rate is a measure of tissue viability.

### Exchange rates are linked to passive permeability, activity, and viability

Fixed tissue is at equilibrium and exchange is entirely passive (*k*_*a*_ = 0). Live tissue maintains a non-equilibrium steady-state under which we assume exchange to be a summation of parallel passive and active components *k* = *k*_*p*_ + *k*_*a*_. Ouabain causes transmembrane ionic gradients to depolarize, bringing the live tissue closer to equilibrium and the overall active transport closer to zero. If we assume *k*_*a*_ to be zero in ouabain-treated spinal cords, after membranes have depolarized, and surface-to-volume ratios to be similar, then we can assess *k*_*p*_ and *k*_*a*_ from comparisons between treatment groups (Fig. 5). We find that fixation increases passive membrane permeability by 240% by comparing fixed to ouabain-treated spinal cords. We find that the exchange rate under normal conditions is approximately 30% passive and 70% active by comparing normal (untreated) to ouabain-treated live spinal cords. OGD of sufficient duration causes energetic failure, akin to an equilibrium state. Exchange rates after 70 min OGD were similar to exchange rates after ouabain treatment. This is likely because both treatments bring the system to a steady-state activity level close to equilibrium.

**Fig. 5.**
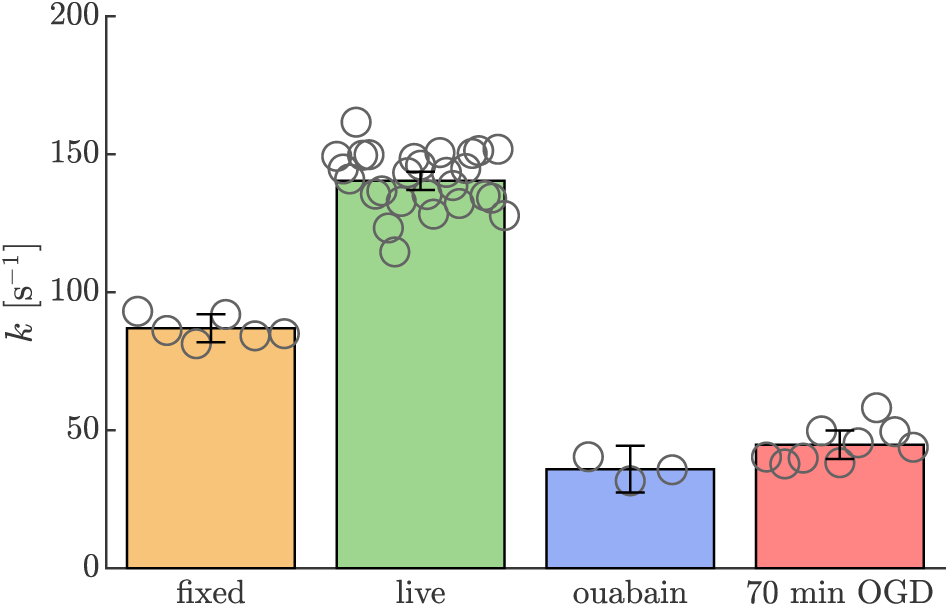
Comparisons between fixed, live (untreated), ouabain-treated, and post-70 min OGD reveal how treatments affect passive and active exchange. Bar graphs present the mean across all measurements (bar height), 95% CI of the mean (whiskers), and mean values from each sample (open circles). Exchange rates were compiled from the 1^st^ 25° C condition in Fig. 1, and from Figs. 3 and 4. The exchange rate of live spinal cords (mean *±* SD *k* = 140 *±* 16 s^*−*1^, *n* = 27) is significantly greater than fixed (*k* = 87 *±* 10s^*−*1^, *n* = 6), ouabain-treated (*k* = 36 *±* 11s^*−*1^, *n* = 3), and spinal cords after 70 minutes of OGD (45*±* 7s^*−*1^, *n* = 9) (*p <* 0.001). Further, the exchange rate of ouabain-treated spinal cords is significantly less than fixed spinal cords (*p <* 0.001). However, exchange rates are not significantly different between ouabain-treated spinal cords and spinal cords after 70 min OGD (*p* = 0.056). Associated data and fits are compared in Fig. S10..

It is important to emphasize that the exchange rate is an absolute, intrinsic measurement. We find across 27 samples that the normative exchange rate lies within a well-defined range: 140 *±* 16s^−1^ (Fig. 5). Similar variability is found from repeated measurements on individual samples, e.g, 153 ±17s^−1^ (Fig. S11). On *n* = 4 samples, exchange rates were recorded for many hours under normal conditions. Three of the samples started with exchange rates within the normal range and showed stability for between 8 and 20 hours. One of the samples started with exchange rates below the normal range and continued to run down. In all cases, when exchange rates fell below the normal range, exchange rates ran down, plateauing at values just above the exchange rates measured on ouabain-treated samples. This further suggests that the exchange rate is a reliable measure of tissue viability.

In contrast, ADC_y_ is a relative measurement (e.g., as a percent change from baseline), not an absolute measurement. Variability of normative ADC_y_ values is greater across samples than within individual samples (0.964 *±* 0.097 *µ*m^2^*/*ms compared to, e.g., 1.017*±* 0.013 *µ*m^2^*/*ms from Figs. S12 and S11 respectively), masking the relative effects of perturbations observed in real-time. We find ADC_y live_ ≈ ADC_y ouabain_ *>* ADC_y 70 min OGD_ *>* ADC_y fixed_ (Fig. S12, unlike Fig. 5). Unlike exchange rates, ADC_y_ values do not differentiate treatment groups based on activity and viability. Instead, results affirm that ADC_y_ values differentiate treatment groups based on shifts in extracellular to intracellular water (52).

## Discussion

Homeostasis, which is the maintenance of a steady-state condition far from equilibrium by the expenditure of metabolic energy, is a defining characteristic of living organisms. We show that the water exchange rate we measure is directly linked to the homeostatic state of parenchymal neural tissue. Moreover, we provide a future direction for interrogating these steady-state processes in the CNS and other living cells or organs.

This study measures intra- and extra-cellular water pools turning over faster than 100 times their volume per second in live CNS tissue. Prior to our finding on *ex vivo* fixed spinal cords (41), there were no reliable measurements of transcy-tolemmal exchange rates this fast in biological systems. Perhaps this is because low-field, high-gradient DEXSY is the first method capable of reliably measuring diffusive exchange in and out of water pools restricted on length scales smaller than a micron (33). Large membrane surface-to-volume (SV) ratios, in combination with high levels of active exchange, leads to the high turnover rates. The neonatal mouse spinal cord consists mostly of gray matter and little myelinated white matter (53, 54). These sub-micron membrane structures are likely glial and neuronal processes (i.e., neurites), which make up 80–90% of the gray matter tissue by volume (55), and may also include organelles (41). Discrepancy from previously reported values, e.g., reports of exchange rates between 1 and 10 s^−1^ (36, 56, 57), can be explained by other methods being sensitive to slowly exchanging water pools associated with larger or less permeable membrane structures, e.g., cell bodies (i.e., soma), and myelinated axons but insensitive to rapidly exchanging pools. Confirming this belief, recent effort to develop models for PFG diffusion MRI of gray matter have found it necessary to account for exchange occurring during the diffusion encoding time, and with rates ≳ 100 s^−1^ (55, 58–61).

The highest exchange rates we measured were 228*±*72 s^−1^ for normal samples at 35°C. Exchange rates could be even greater at 37°C *in vivo* as this is the most viable condition for the tissue. We found that viability was difficult to maintain at 35°C. For this reason, elecrophysiologists typically study the neonatal mouse spinal cord at room temperature around 25°C (62). Live samples still showed signs of activity at 7°C — in particular, fixed and live spinal cords had similar exchange rates even though fixation increases passive permeability. In contrast, Esmann & Skou measured activity of Na^+^*/*K^+^–ATPase isolated from ox brain to be near zero at 7°C (63). This discrepancy warrants further research into the mechanisms of active water transport.

Active water exchange was found to be linked to ion transport (Fig. 5). One possible mechanism is that water cotransports with ions directly through Na^+^*/*K^+^–ATPase or other transport proteins. In this way, a number of transporters including KCC4 and NKCC1 have been shown to cotransport water against an osmotic gradient (12). Another similar mechanism is that ion transport creates a local osmotic gradient which drives water transport through nearby water channels (e.g., aquaporin), as shown for SGLT1 and EAAT1 (12). Both mechanisms transport up to hundreds of water molecules per ion or metabolite. Regardless of the actual mechanism, active water exchange was found to account for at least 70% of the total exchange rate, leaving only 30% as passive.

Most studies of water transport in biological systems consider it to be solely passive (48, 64). Activation energies (*E*_*a*_) of passive water permeability are independent of membrane SV ratio and are therefore often used for comparison (48). These studies show *E*_*a*_ values trend with lipid bilayer composition and concentration of passive water channels such as aquaporins (48, 64). Lipid bilayers have *E*_*a*_ between 33 to 40 kJ*/*mol because permeability depends on membrane fluidity which varies strongly with temperature. Aquaporins increase permeability but reduce *E*_*a*_ towards the *E*_*a*_ for self-diffusion of water (*E*_*a*_ = 18 −20 kJ*/*mol (47)). Exemplifying these extremes, *E*_*a*_ = 40 kJ*/*mol was found for Baker’s Yeast in which aquaporin channels were presumed to be closed (34), and *E*_*a*_ values near 25 kJ*/*mol were consistently reported for red blood cells (65) in which aquaporins are highly expressed (66, 67). We found *E*_*a fixed*_ = 21*±*8 kJ*/*mol, consistent with permeability by diffusion through pores opened during fixation (discussed below).

For passive water permeability, lower *E*_*a*_ values are associated with higher permeability. Active water transport alters this trend because enzymatic activity increases strongly with temperature. Isolated Na^+^*/*K^+^–ATPase (63) and water cotransport (12) both have *E*_*a*_ ∼ 100 kJ*/*mol at physiological temperatures (63). We found *E*_*a live*_ *> E*_*a fixed*_ and *k*_*live*_ *> k*_*fixed*_ indicating active water exchange exists. Values for *E*_*a live*_ = 36*±*7 kJ*/*mol are between values for active water cotransport and passive water self-diffusion, consistent with exchange in live CNS tissue being both active and passive.

The *E*_*a*_ of ADC_*y*_ in fixed and live spinal cords are less than values for pure aCSF and reported values for water (47). This is because the temperature dependence is dominated by hindered diffusion within the tissue. As the length scale of water diffusion increases with temperature, water experiences more interactions with membranes, causing the temperature dependence of diffusion to be less than that of pure water. The *E*_*a*_ of ADC_*y*_ was found to be not significantly different between live and fixed spinal cords. This indicates that ADC is not directly sensitive to cellular activity, supporting evidence (68, 69) which discredits the proposition that it directly detects neuronal activation (the premise of diffusion fMRI) (70).

While we compare fixed vs. live tissue primarily to test whether NMR properties are linked to activity, the data also provides information about the effects of fixation. Shepherd *et al*. performed a similar study on fixed and perfused rat brain cortical slices (71). However, their methods involved stopping circulation during measurements, which could have affected tissue viability. We therefore compare our findings with theirs, under the plausible assumption that their perfused tissue had limited viability and active exchange. We found fixation increased the passive exchange rate *k*_*p*_ by 240% (Fig. 5), strikingly similar to 239% reported by Shepherd *et al*. (71). We also found that fixation opened pores in the membrane large enough for sucrose molecules to penetrate (see Figs. S6, S7, and S8). Together, these findings indicate that water permeability in fixed tissue is determined primarily by diffusion through pores opened during fixation. We found fixation decreased ADC_*y*_ by 20%. Fixation shrinks the extracellular space from 20% to 5% (71), which reduces the fraction of more mobile extracellular water and increases the fraction of less mobile intracellular water. This re-partitioning of water during fixation reduces ADC_*y*_ overall. Shepherd *et al*. reported that diffusivity *increased* upon fixation, however this discrepancy could be related to the longer diffusion encoding times used in their study (10 to 50 ms vs. 0.2 to 0.5 ms for our ADC_*y*_ measurements), or to the perfused sample conditions. We found that fixation caused a 15% increase in *R*_1 DW_ at 0.32 T. *R*_1 DW_ for fixed spinal cords is higher than for live spinal cords because formaldehyde crosslinks reduce the rotational mobility of proteins, which causes water to relax faster. While *R*_1_ is field strength-dependent and we report a diffusion-weighted *R*_1_, our results are consistent with Shepherd *et al*. who report that fixation increased *R*_1_ by 21% at 17.6 T (71). These effects of fixation suggest that fixed tissue may not always be an appropriate sample when developing MRI methods intended for *in vivo* applications. Alternatively, we show viable CNS tissue is a useful model for studying how pathologies affect MR parameters without *in vivo* confounds.

Diffusion-weighted imaging (DWI) is the gold-standard for identifying stroke because the ADC decreases significantly in affected areas minutes after stroke due to cytotoxic edema (72, 73). However, the diffusion coefficient alone cannot differentiate recoverable tissue from permanently damaged tissue (74–76). DWI shows similarly reduced diffusivities throughout an ischemic area, masking the heterogeneous effects on tissue metabolism (77, 78). The ADC of a lesion begins to normalize during the first few days following a stroke, sometimes indicating tissue recovery (73). However, in many cases that the tissue is actually damaged, the ADC will appear to recover or “pseudo-normalize” and will even increase to values higher than the surrounding normal tissue due to necrosis and loss of membrane integrity (73, 79, 80). The timecourse of ADC_*y*_ during OGD recapitulates *in vivo* findings — it decreases initially upon switching to OGD, and pseudo-normalizes and sometimes overshoots baseline values during recovery (Figs. 4 and S9) — thus confirming that ADC is not specific to tissue damage.

In contrast, timecourses of exchange rates during OGD (Fig. 4), stability tests (Fig. S11), and rundown show trends consistent with the exchange rate measuring viability. A potential mechanism is that the water exchange rate is linked to the cellular homeostatic state. The exchange rate is highly regulated, stable, and absolute because cellular processes compensate to maintain the homeostatic state. The exchange rate drops when homeostasis is lost and viability is compromised.

Further research is needed to translate low-field, high-gradient DEXSY to *in vivo* human brain MRI. Future directions could broadly involve development of static gradient methods or PFG methods. However, either direction may require compromising gradient strength, which with current technology is limited by the maximum bandwidth of radiofrequency (RF) probes (needed to excite a sizable volume of protons under a static gradient) or by the maximum gradient strength of PFG coils and amplifiers, as well as by biological constraints such as RF heating and peripheral nerve stimulation. One major question is whether weaker gradients and longer diffusion encoding times will provide the same sensitivity to active exchange found in this study.

We establish the exchange rate as an absolute, intrinsic measure of cellular function and viability, and our findings lay the groundwork for developing the exchange rate as a potential imaging biomarker more directly linked to cellular metabolic activity than current functional MRI (e.g., BOLD fMRI) and to permanent tissue damage than current structural MRI (e.g., diffusion MRI).

## Methods

Details of the methods used here can be found in Refs. (41, 42) and are described briefly below.

### Ethics statement for animal experimentation

All experiments were carried out in compliance with the *Eunice Kennedy Shriver* National Institute of Child Health and Human Development Animal Care and Use Committee (Animal Study Proposal (ASP) # 21-025) and the National Institute of Neurological Disorders and Stroke Animal Care and Use Committee (ASP # 1267–18).

### Sample preparation and materials

All experiments were performed on Swiss Webster wild type (Taconic Biosciences, Rensselaer, NY, USA) mice between postnatal day 1 to 4. The mouse spinal cords were isolated and placed in a dissecting chamber perfused with cold low-calcium high magnesium artificial cerebrospinal fluid (aCSF, concentrations in mM: 128.35 NaCl, 4 KCl, 0.5 CaCl_2_ ·2H_2_O, 6 MgSO_4_ · 7H_2_O, 0.58 NaH_2_PO_4_ ·H_2_O, 21 NaHCO_3_, 30 D-glucose) bubbled with 95% O_2_ and 5% CO_2_). Live spinal cords were transported in low-calcium, high-magnesium aCSF to the NMR experimental apparatus. Fixed spinal cords were kept overnight in 4% paraformaldehyde at 4°C and then stored in phosphate buffered saline (PBS) at 4°C, and washed with aCSF three times over the course of two days to remove any residual paraformaldehyde prior to experiments. Experiments on fixed and live spinal cords were performed using normal aCSF (same concentrations as the low-calcium high-magnesium solution except for 1.5 mM CaCl_2_ · 2H_2_O and 1 mM MgSO_4_ · 7H_2_O.

### Hardware, setup, and experimental conditions

A PM-10 NMR MOUSE single-sided permanent magnet (81) (Magritek, Aachen Germany) and a Kea 2 spectrometer (Magritek, Wellington, New Zealand) were used to perform NMR experiments at B_0_ = 0.3239 T (proton *ω*_0_ = 13.79 MHz) and *g* = 15.3 T*/*m. A test chamber and RF probe were fabricated to maximize SNR and maintain live spinal cord viability. A diagram of the experimental setup is shown in Fig. 6.

**Fig. 6.**
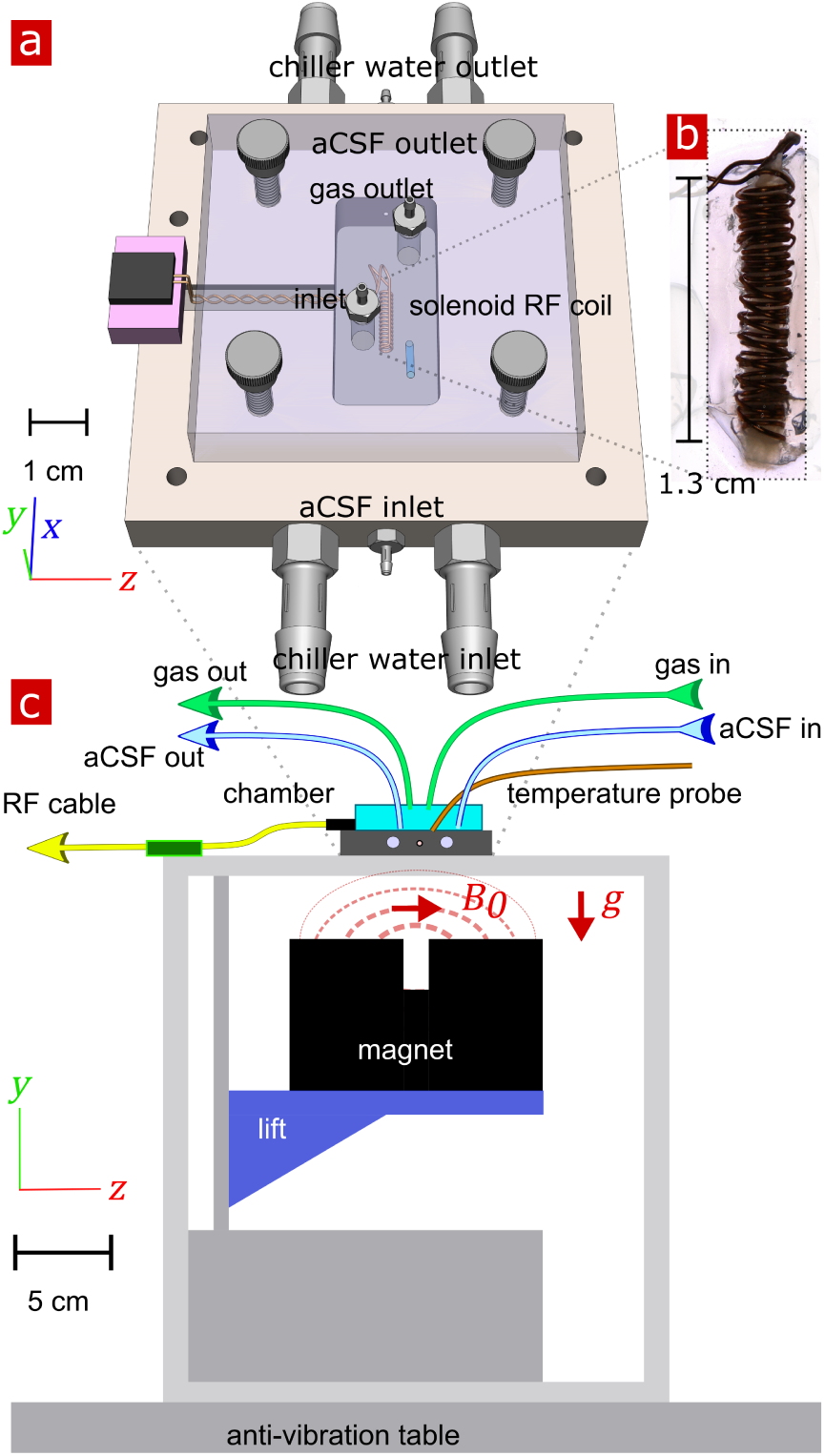
Experimental setup. (a) 3-D technical drawing of the test chamber. aCSF inlet/outlets and temperature probes are omitted for simplicity. (b) Image of the solenoid RF coil containing a mouse spinal cord. (c) Technical drawing of the experimental setup. Vectors B_1_, *g* and B_0_ point in the *x, y*, and *z* directions, respectively

Sample temperature was controlled by heat exchange with one of two circulating water baths (Accel 250 LC, Thermo Scientific, USA, and WCR-P6 Precision Regulated Bath Circulator, Daihan Scientific, Lennox Laboratory supplies, Ireland). The chamber was made in-house out of aluminum to facilitate heat transfer with the water circulating through channels cut through the chamber. Three-way valves located upstream and downstream of the chamber were used to rapidly switch between water baths at different temperatures. While one chiller maintains the sample temperature at a particular set point, the temperature of the other chiller can be changed and slowly equilibrated. Sample temperature was monitored by a fiber optic sensor (PicoM, Opsens Solutions Inc., Québec, Canada). After switching temperatures via the three-way valves, the sample temperature equilibrated in roughly 5 minutes and then remained stable ± 0.5°C there-after. (See representative temperature log, Fig. S1.) NMR measurements were started 10 minutes after the sample temperature stabilized. The temperature was set to 25° C for experiments where temperature was not varied.

The same experimental conditions were used for live and fixed spinal cords. Humid 95% O_2_ and 5% CO_2_ gas flowed into the top of the sealed chamber. aCSF was bubbled with 95% O_2_ and 5% CO_2_. A peristaltic pump circulated aCSF media continuously through the chamber at 2 ml/min.

Oxygen glucose deprivation (OGD) studies involved switching the media inflow between normal aCSF to glucose-free aCSF (made with 30 mM sucrose to keep osmolarity constant) bubbled with 1% O_2_, 5% CO_2_, and 94% N_2_. At the same time, gas flowing into the top of the sealed chamber was switched to humid 1% O_2_, 5% CO_2_, and 94% N_2_.

### NMR methods

Experimental protocols involved looping through sets of diffusion experiments and rapid DEXSY experiments to acquire repetitions of each. Noise and RF probe tuning were monitored at the beginning of each loop. Experiments used repetition time (TR) = 2 s, 2 *µ*s 90°/180° hard RF pulses with amplitudes = −22/−16 dB. Carr–Purcell– Meiboom–Gill (CPMG) acquisition blocks used 2000 or 8000 echoes with 25 *µ*s echo time, 4 *µ*s acquisition time and 0.5 *µ*s dwell time (82, 83). The static gradient was in the *y* direction (Fig. 6) and defined the slice and diffusion encoding directions. Signal was acquired from approximately a 400 *µ*m slice through the length of the spinal cord.

Diffusion experiments were performed using a standard sequence involving a spin echo (SE) for diffusion encoding followed by a CPMG signal acquisition (84). *τ* (defined as half the SE echo time) was varied linearly from 0.05 to 3.3 ms over 22 steps with 4 scans per *τ*. This corresponds to *b*-values ranging from 0.001 to 400 ms*/µ*m^2^ where *b* = 2*/*3*γ*^2^*g*^2^*τ* ^3^ (82, 85).

Points two through four (*τ* = 0.2048 to 0.5143 ms, *b* = 0.096 to 1.5 ms*/µ*m^2^) of diffusion data were fit with *I*(*b*) = *I*_0_ exp(−*b* ADC) to estimate the Apparent Diffusion Coefficient and *I*_0_. For measurements on spinal cords, the term ADC_*y*_ is used, acknowledging that diffusion may be anisotropic but was measured only in the *y* direction, perpendicular to the cord.

Points 12 through 22 were fit with a model for diffusion within cylindrical restrictions oriented perpendicular to the gradient direction with a constant gradient (86, 87) and incorporating exchange (88),

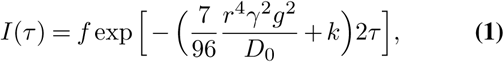

to estimate the restriction radius *r* and the restricted fraction *f*. Each estimate incorporated *k* measured from the rapid exchange experiment during the same set.

Rapid exchange experiments were performed using a DEXSY sequence involving two spin echoes separated by a mixing time *t*_*m*_ and a CPMG acquisition (41) and following Method 3 in Ref. (42). The sequence used 8 phase cycle steps to avoid unwanted coherence transfer pathways (41). For experiments involving temperature perturbations, DEXSY data points were acquired with (*τ*_1_, *τ*_2_) combinations (0.200, 0.213), (0.200, 0.735), (0.593, 0.580), and (0.735, 0.200) ms, corresponding to (*b*_1_, *b*_2_) = (0.089, 0.1080), (0.089, 4.417), (2.320, 2.170), and (4.417, 0.089) ms*/µ*m^2^. For all other experiments, combinations (0.200, 0.213) and (0.735, 0.200) were omitted due to the findings from Ref. (42) that these points are redundant. Each (*τ*_1_, *τ*_2_) combination was acquired with 8 scans and with *t*_*m*_ = [0.2, 1, 2, 4, 7, 10, 20, 40, 80, 160, 300] ms. The signal from (*τ*_1_, *τ*_2_) = (0.200, 0.735) and (0.735, 0.200) was averaged and fit with *I*(*t*_*m*_) = *I*_0_ exp(*−t*_*m*_*R*_1 DW_) to estimate *R*_1 DW_. The same signal was also fit with 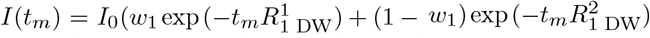. The resulting model was subtracted from the signal from (*τ*_1_, *τ*_2_) =(0.593, 0.580) and the signal was fit with *I*(*t*_*m*_) = *I*_0_ exp(− *t*_*m*_*k*) + *B* to estimate *k*. Representative data and fits are shown in Fig. S13.

Saturation recovery experiments were performed during the OGD study to measure *R*_1_ as a means of monitoring pO_2_ changes. Experiments used 6 recovery times exponentially spaced between 0.067 and 6 ms. Sensitivity of *R*_1_ to pO_2_ was confirmed with measurements on pure aCSF, circulating through the chamber and bubbled with 1% or 95% O_2_ gas, and was also used to determine the gas flow/bubbling rate sufficient to reach O_2_ saturation.

An Arrhenius model of the form *fn*(*T* ^*−*1^) = *A*exp(−*E*_*a*_ */ RT*), where *R* = 8.3145 ×10^−3^ kJ*/*(mol K) is the ideal gas constant, was fit to measurements of *k, T*_1_, or ADC_*y*_ as a function of the inverse of the absolute temperature *T* ^*−*1^ to estimate activation energies (*E*_*a*_) associated with each metric for each sample.

## Statistical methods

Data was analyzed using MATLAB R2020a. Experimental results involving many measurements are presented as box and whisker plots and violin plots in order to provide a full sense of the structure and variability of the measured data. Box plots show the median (red middle line), the 25^th^ percentile (bottom line), and the 75^th^ percentile (top line) (89). Notches in the box plot show the 95% confidence interval (CI) of the median. (Note that the notches sometimes extend further than the 25^th^ or 75^th^ percentiles.) Violin plots show a smooth probability density function (pdf) for the distribution of measured values (90). Means, standard deviations, and 95% CIs are used in other cases, as noted in figure captions. Both 95% CIs of the median and two-sample (unpaired) t-tests assuming equal variance (*α* = 0.05) are used for hypothesis testing between sample groups. Pearson correlations were analyzed using the MATLAB corrcoef function to estimate correlation coefficients and *p*-values. Cross-correlations were analyzed using the MATLAB xcorr function to estimate the time lag between effects in simultaneously acquired real-time data.

## Data availability statement

All data and code used in analysis will be made publicly available as supplementary material upon publication.

## ACKNOWLEDGEMENTS

We thank Dave Ide for machining specialty parts and Marcial Garmendia-Cedillos for soldering and creating the technical drawing (Fig. 6), Randall Pursley, Helmut Merkle and Tom Pohida for electrical engineering advice, and Alexandru Avram, Dan Benjamini, Michal Komlosh, and Samuel Neuman for helpful discussions. NHW was funded by the NIGMS PRAT Fellowship Award # FI2GM133445-01. RR, TXC, and PJB were supported by the IRP of the NICHD, NIH. MF and MJO were funded by the IRP of the NINDS.

## Supplementary figures

**Fig. S1.**
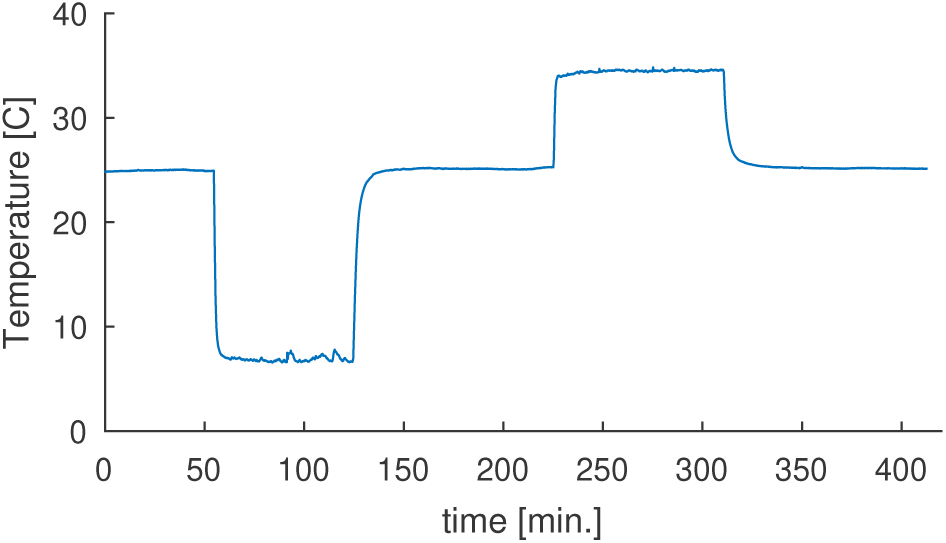
Representative log of sample temperature during an experiment. Experimental protocol varied temperature from 25–7– 25–35–25°. Temperature was recorded in the chamber using a fiber optic sensor (see Methods).

**Fig. S2.**
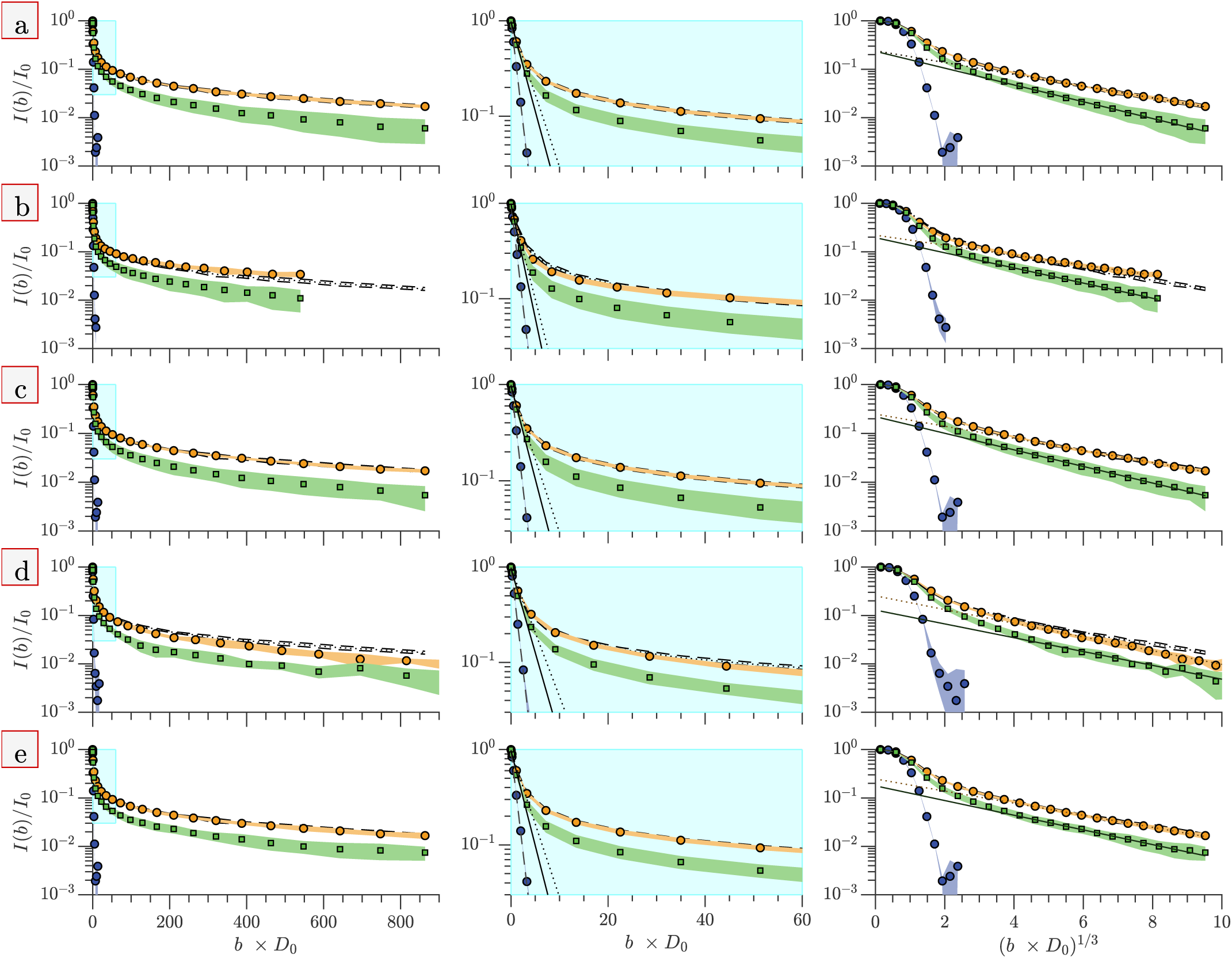
Diffusion signal comparison. Diffusion signal intensity from measurements on *n* = 6 fixed (orange circles) and *n* = 7 live (green squares) spinal cords, and aCSF (purple circles) performed first at 25°C (row a), followed by 7°C (b), 25°C (c), 35°C (d), and 25°C (e). Means (symbols) and standard deviations (shaded bands) are presented. The three columns show the same data plotted differently. Figures in column 1 are plotted as a function of *b × D*_0_. On this scale, freely diffusing water decays exponentially. Column 2 figures zoom in on the initial signal attenuation, corresponding to a region shaded light blue in the column 1 figures. The straight dotted, solid, and dashed lines are best fits of the initial decay, where the slopes correspond to ADC_*y*_. Figures in column 3 are plotted as a function of (*b* × *D*_0_)^1*/*3^. On this scale, restricted water decays roughly exponentially. The straight dotted and solid lines are fits of the final decay at long *τ* (large *b*^1*/*3^) (points 12–22). Lines extrapolate back to *b*^1*/*3^ = 0, corresponding to the restricted volume fraction *f*. In all figures, the mean (dotted line) and *±* standard deviation (dashed lines) of the fixed samples from the first 25°C experiments are re-plotted to show how the signals collapse with the non-dimensionalization and to show how well the signals recover when returning back to 25°. The variable *b* is non-dimensionalized by the measured diffusion coefficients of aCSF at 7°, 25°, and 35°C, *D*_0_ = 1.35, 2.16 and 2.74 ms*/µ*m^2^.

**Fig. S3.**
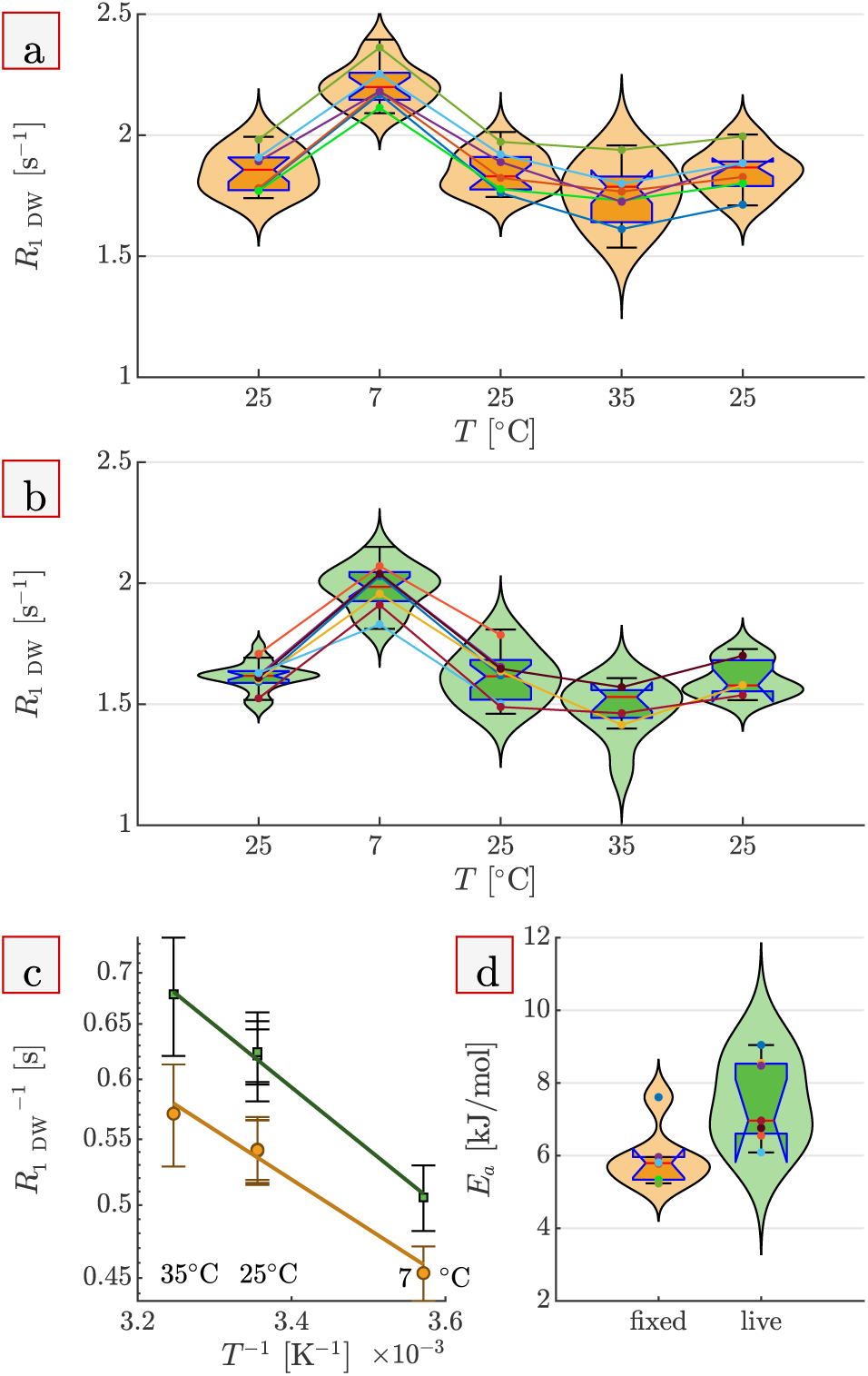
Temperature dependence of diffusion-weighted spin-lattice relaxation. Diffusion-weighted spin lattice relaxation rate *R*_1_ of fixed (a) and live (b) spinal cords at each temperature condition. (c) Arrhenius plot of *R*_1_^−1^ (≈ *T*_1_) for fixed (orange) and live (green) spinal cords. (d) Boxplots comparing activation energies between fixed and live samples. A greater description of plot details is provided in the caption of Fig. 1. 95% CI of median values indicate that the activation energy of *R*_1_^−1^ is similar between live and fixed spinal cord. t tests indicate activation energy of *R*_1_^−1^ is significantly greater for live spinal cords (*M* = 7.5 kJ*/*mol, *SD* = 1.2 kJ*/*mol) compared to fixed spinal cords (*M* = 6.0 kJ*/*mol, *SD* = 0.86 kJ*/*mol), *t*(11) = 2.7, *p* = 0.02. Activation energy values for *R*_1_^−1^ and ADC_*y*_ (Fig. 2 d) are similar. This indicates that *R*_1_ and ADC_*y*_ are similarly affected by tissue microstructure.

**Fig. S4.**
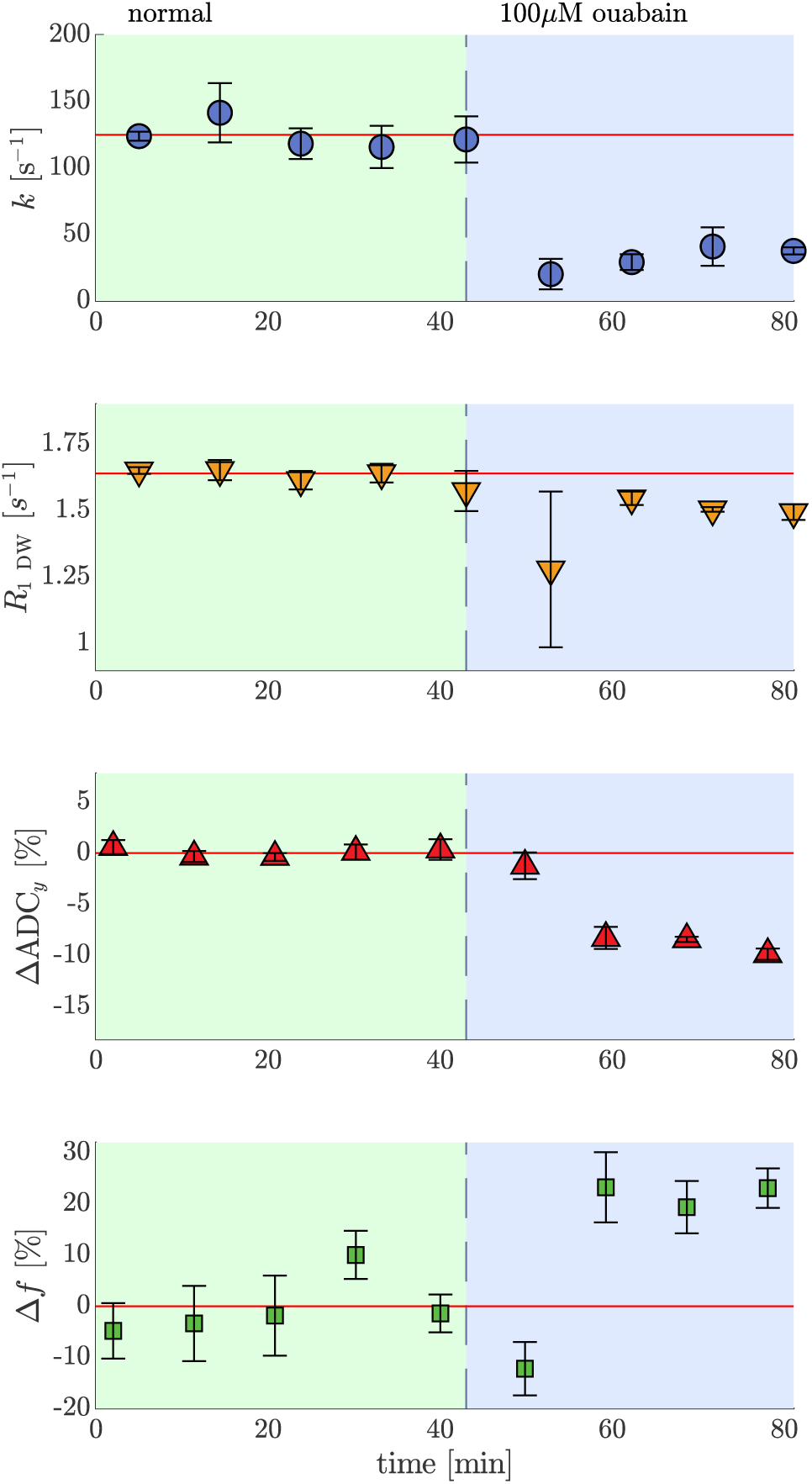
Effect of 100*µ*M ouabain on NMR properties. Real-time exchange rate, diffusion weighted spin-lattice relaxation rate, apparent diffusion coefficient, and restricted volume fraction (from top to bottom) measurements on live spinal cords under normal condition (green) and after (magenta) addition of 100 *µ*M ouabain. Mean values (circles) and standard deviations (whiskers) from n=3 samples are presented. *R*_1 DW_ decreased by 8 *±* 3%. *f* increased by 22 *±* 6 %.

**Fig. S5.**
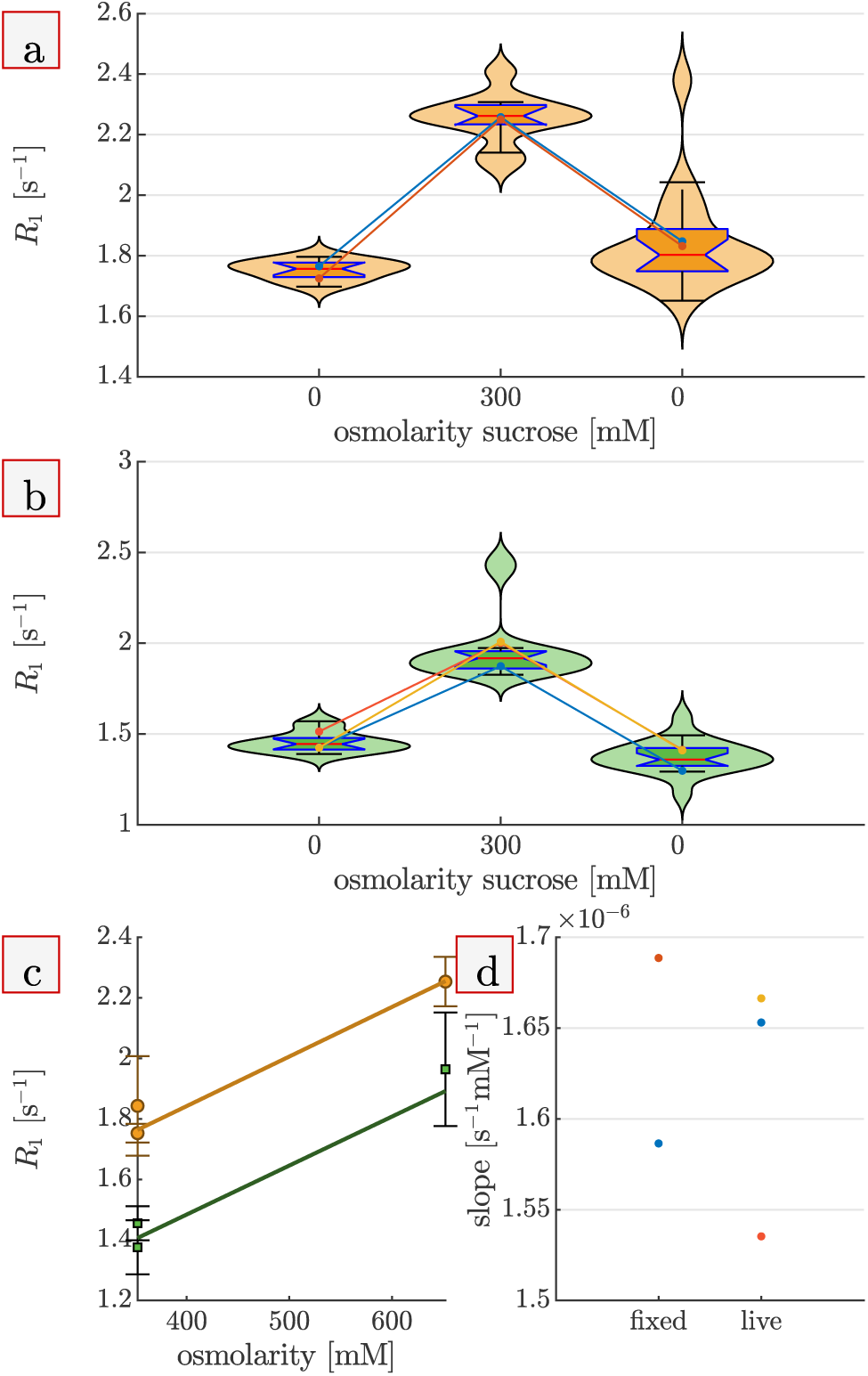
Osmolarity dependence of *R*_1_. Diffusion weighted *R*_1_ for *n* = 2 fixed (a) and *n* = 3 live (b) samples during osmolarity perturbation with sucrose. (c) Plots of *R*_1_ as a function of osmolarity. (d) The slopes from (c) for each sample. Slopes are positive and similar for live and fixed, indicating that sucrose penetrates into the tissue similarly for live and fixed samples.

**Fig. S6.**
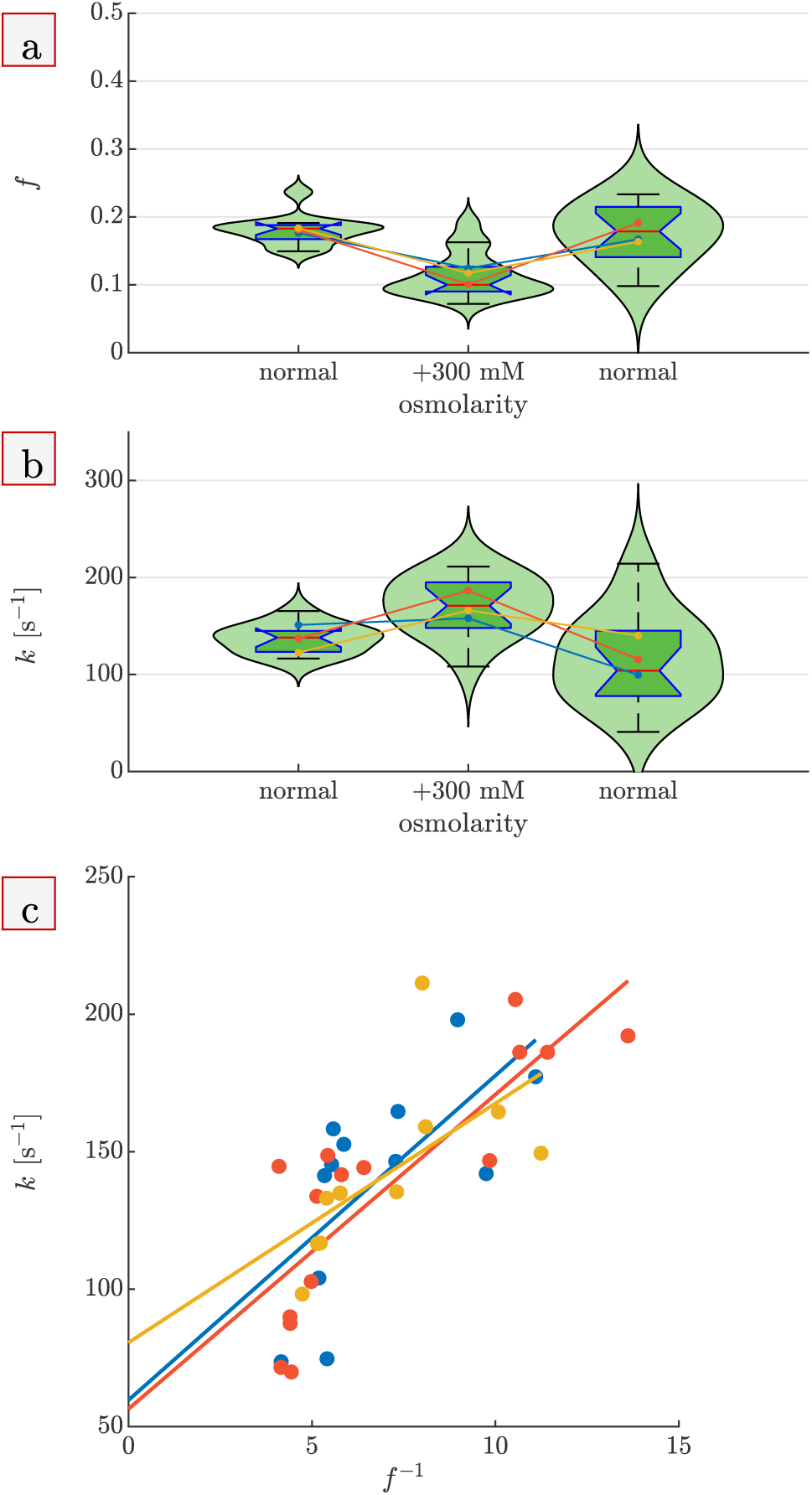
Correlation between *f* ^−1^ and *k*. Measurements of (a) restricted fraction (*f*) and (b) exchange rate (*k*) during osmotic perturbations from normal aCSF to aCSF +300mM sucrose and back on *n* = 3 live spinal cords. c) Values of *f* ^−1^ and *k* acquired from the same set are plotted for each sample and show positive correlations (*p* = 0.019, 0.0001, and 0.039 with correlation coefficients 0.66, 0.83, and 0.63). A correlation is expected through the linear dependence of *k* on the membrane surface-to-volume (SV) ratio. This result is consistent with DEXSY measuring water exchange between the intra- and extracellular space.

**Fig. S7.**
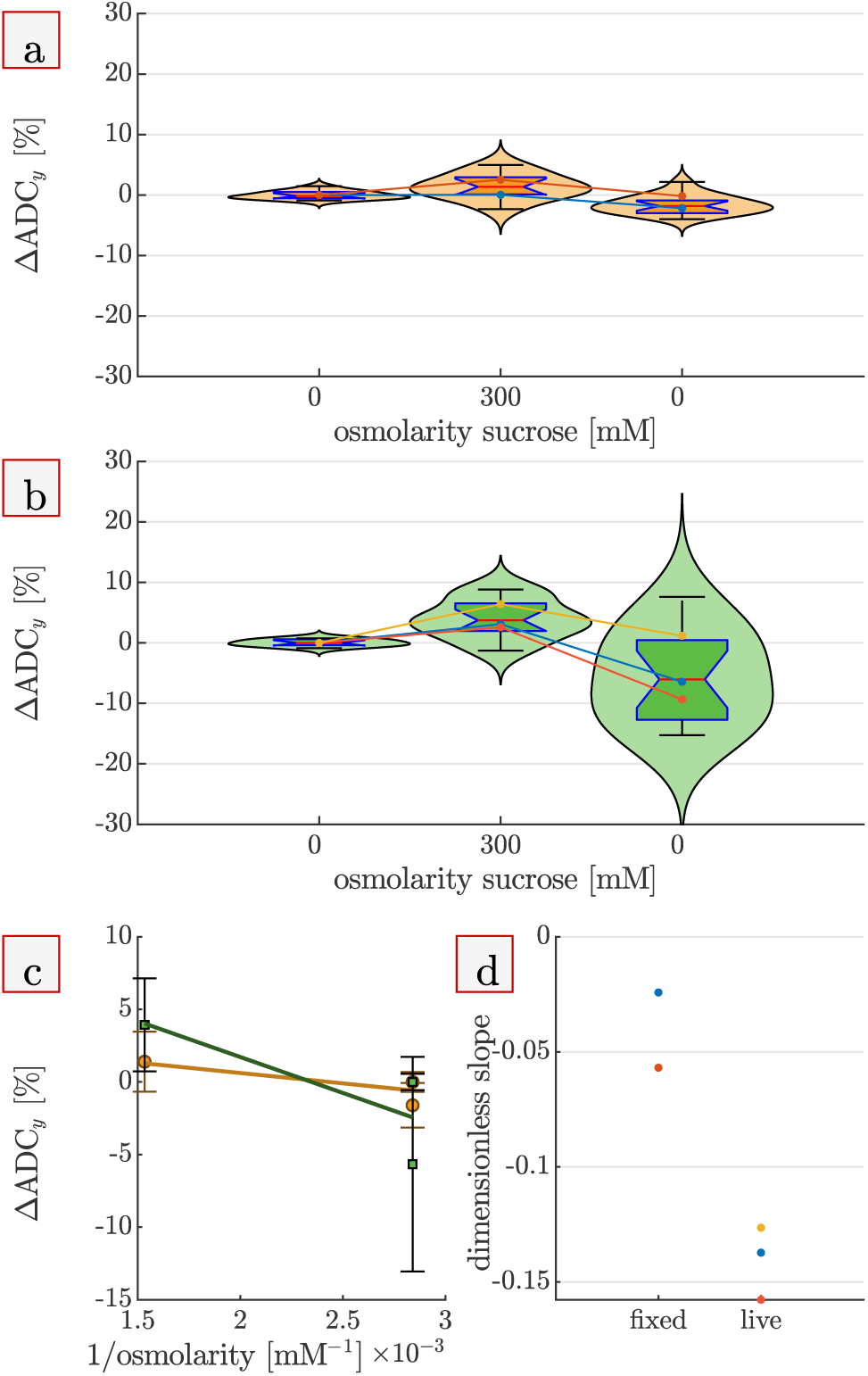
Osmolarity dependence of ADC_*y*_. Percent change in ADC_*y*_ for fixed (a) and live (b) samples during osmolarity perturbation with sucrose. Boxplots show that +300 mM sucrose increases ADC_*y*_ significantly in live samples, consistent with water re-partitioning from the intracellular space to the extracellular space, increasing the fraction of more mobile extracellular water, decreasing the fraction of less mobile intracellular water ADC_*y*_ increases negligibly in fixed samples, consistent with sucrose penetrating through pores in cell membranes opened during fixation and hence having no osmotic effect. In live samples, ADC_*y*_ decreases below baseline when washing back to normal media with +0 mM sucrose, perhaps due to cellular swelling. (c) Plots of ADC_*y*_ as a function of osmolarity. (d) The slopes from (c) for each sample.

**Fig. S8.**
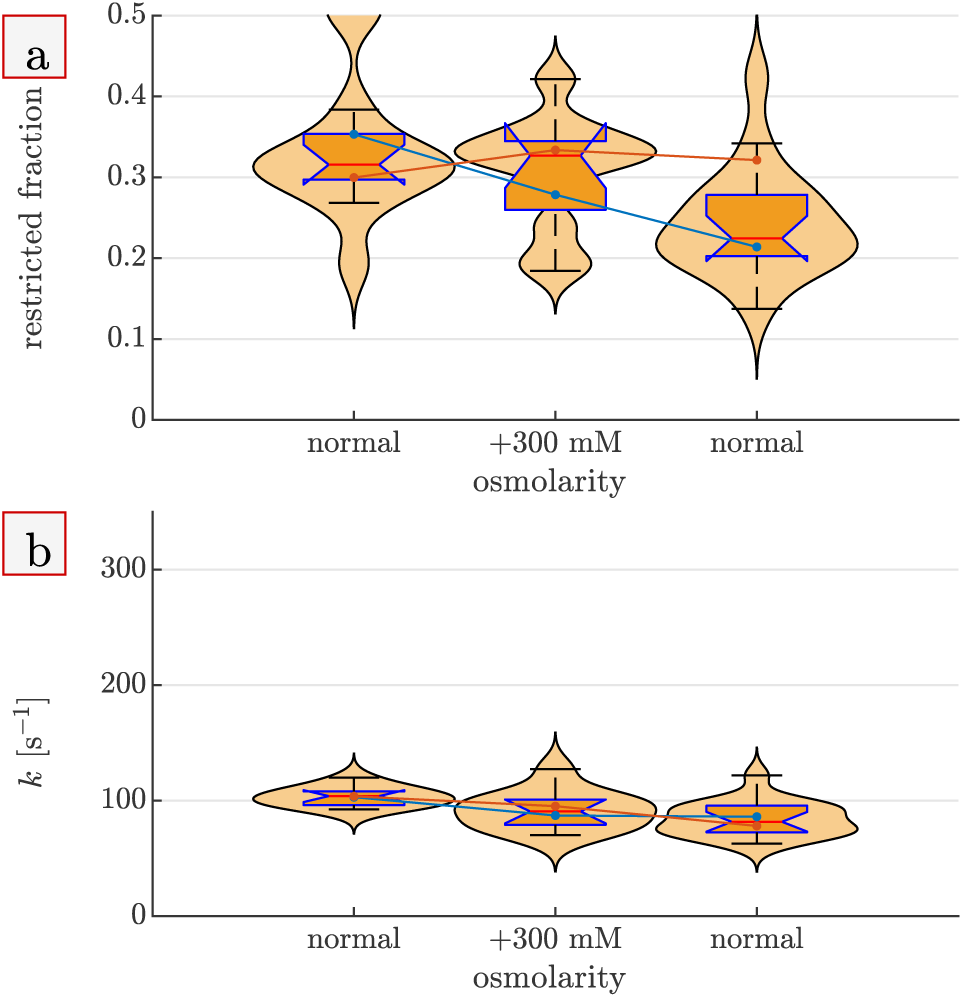
Osmolarity dependence of *f* and *k* in fixed spinal cords. The (a) restricted fraction and (b) exchange rate from *n* = 2 fixed samples undergoing perturbations from normal aCSF to aCSF with 300 mM sucrose back to normal. Unlike for live samples, the change in restricted fraction is insignificant between normal and +300 mM sucrose, indicating that sucrose does not act as an osmolyte on fixed samples; membranes become permeable to sucrose during fixation. There is no change in *k* between normal and +300 mM sucrose because the volume and permeability are unaffected in this perturbation on fixed samples.

**Fig. S9.**
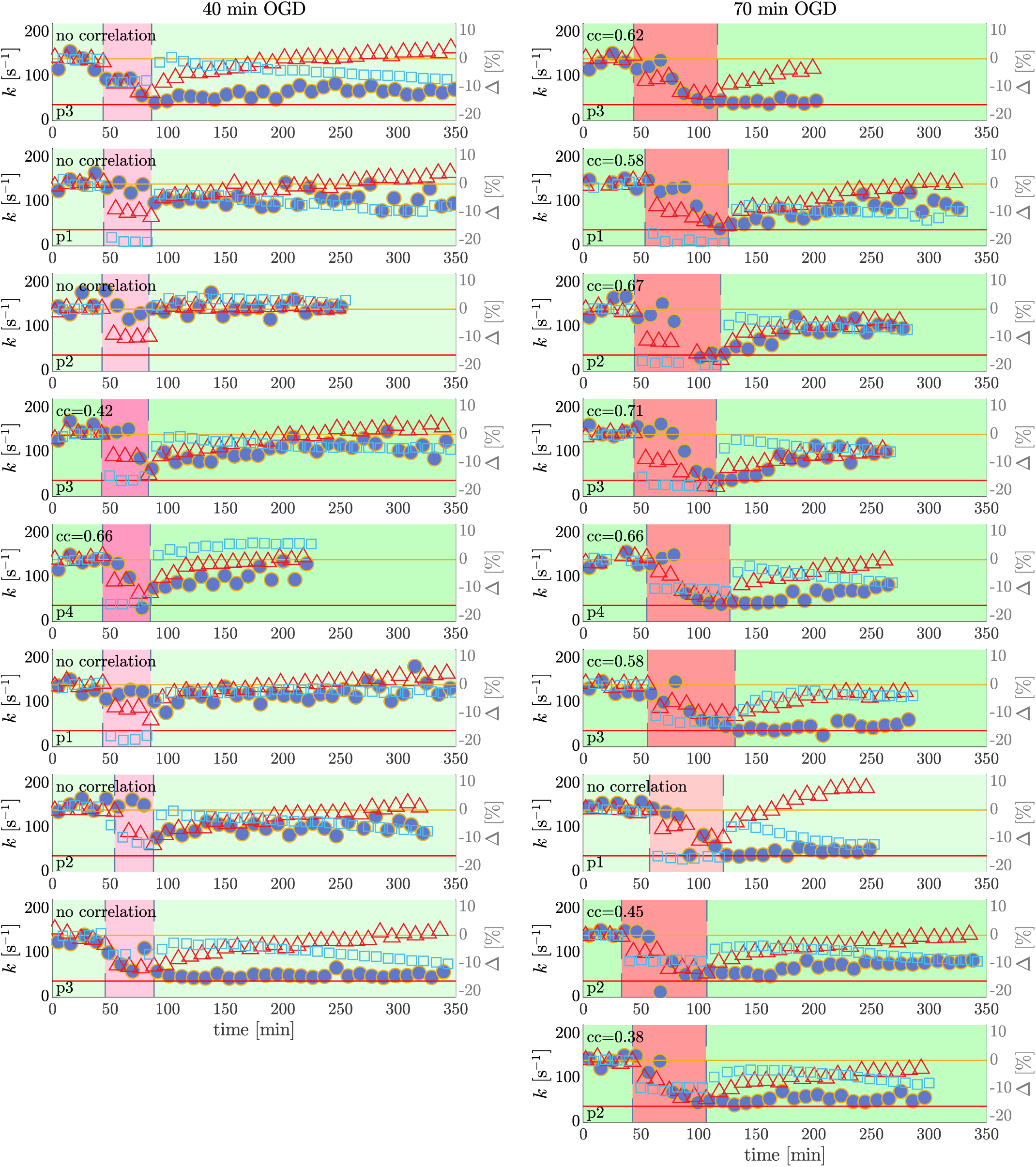
Timecourses for all samples undergoing OGD. Timecourse of exchange rate, apparent diffusion coefficient, and spin lattice relaxation rate for all samples undergoing 40 min (left column, n=9) and 70 min (right column, n=9) of oxygen glucose deprivation (OGD, 1% pO_2_, 0 mM glucose). Exchange rate (solid blue circles) values can be read from the left-hand y-axis. Diffusion coefficients (open red triangles) and spin-lattice relaxation rates *R*_1_ from saturation recovery measurements (open blue squares) are presented as percent change from the initial baseline and values can be read from the right-hand y-axes. Postnatal day of spinal cord dissection (P1, p2, p3 or p4) is shown in the bottom left-hand corner of each plot. Pearson correlation coefficients (*cc*) and associated *p* values for the comparison between exchange rate and diffusion timeseries were calculated. Of all 18 samples, 8 samples did not show a significant correlation between diffusion and exchange (*p >* 0.05 and *cc <* 0.2). The backgrounds for the plots of these samples are shaded lighter. The background are shaded darker and *cc* values are presented in the upper left corner for each plot of the 10 samples which did have a significant correlation (*p <* 0.05). These samples had a (mean*±*SD) *cc* = 0.58 *±* 0.12.

**Fig. S10.**
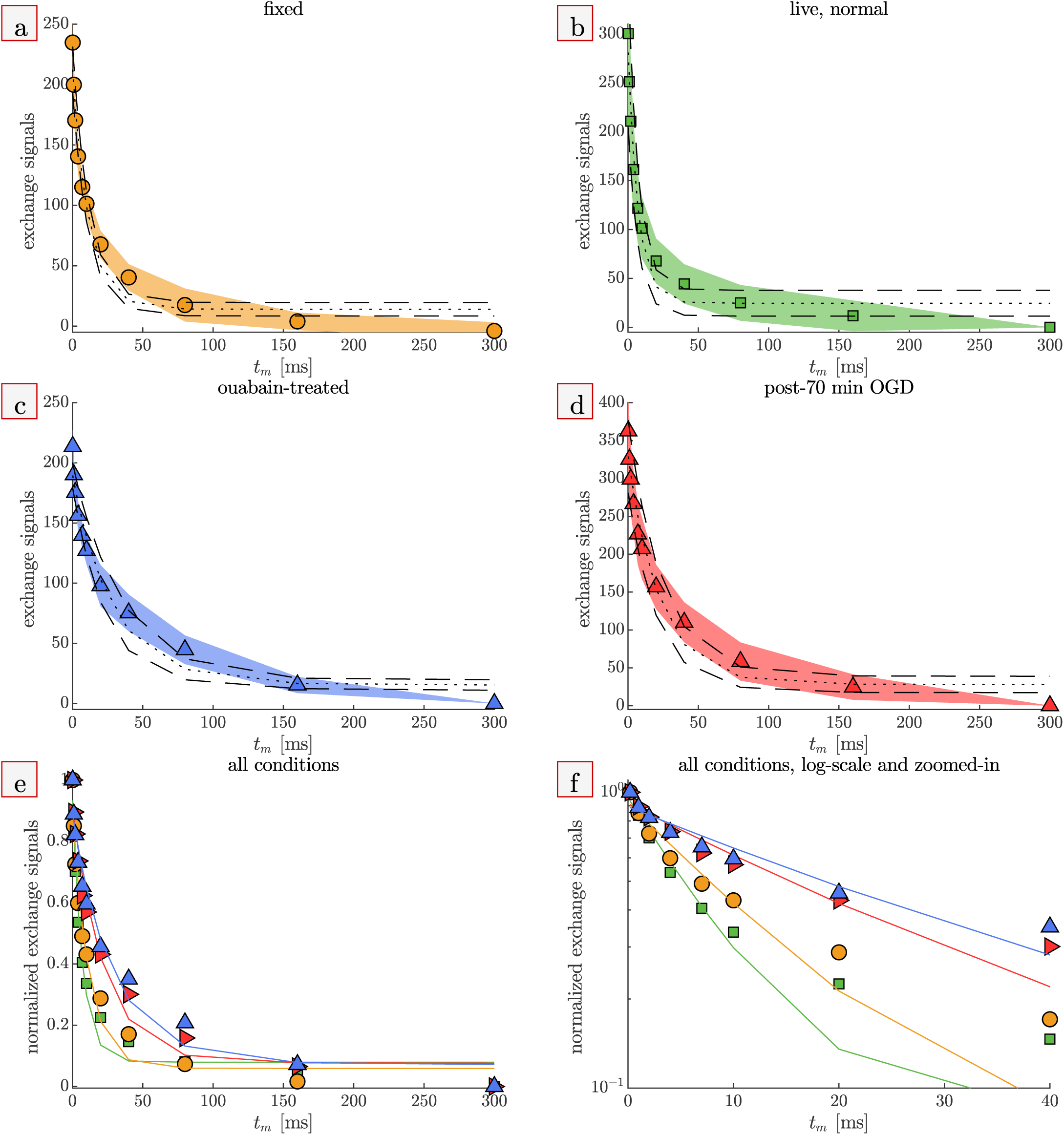
Exchange data and fits compared between fixed, live (untreated), ouabain-treated, and post-70 min OGD. a-d) Mean (symbols) and standard deviations (shaded bands) of exchange signals, and mean (dotted lines) and standard deviations (dashed lines) of exchange rate model fits. Data was compiled from the 1^st^ 25°C condition in Fig. 1 and from Figs. 3 and 4. Exchange rates are compared in Fig. 5. e) Mean of data (symbols) and fits (lines) compared between treatment groups. f) Comparison of the initial (*t*_*m*_ = 0 *−* 40 ms) decay on a log-y axis. Data shows multiexponential behavior, with a faster initial decay and a slower final decay relative to the single exponential fits. See Methods and Fig. S13 for details.

**Fig. S11.**
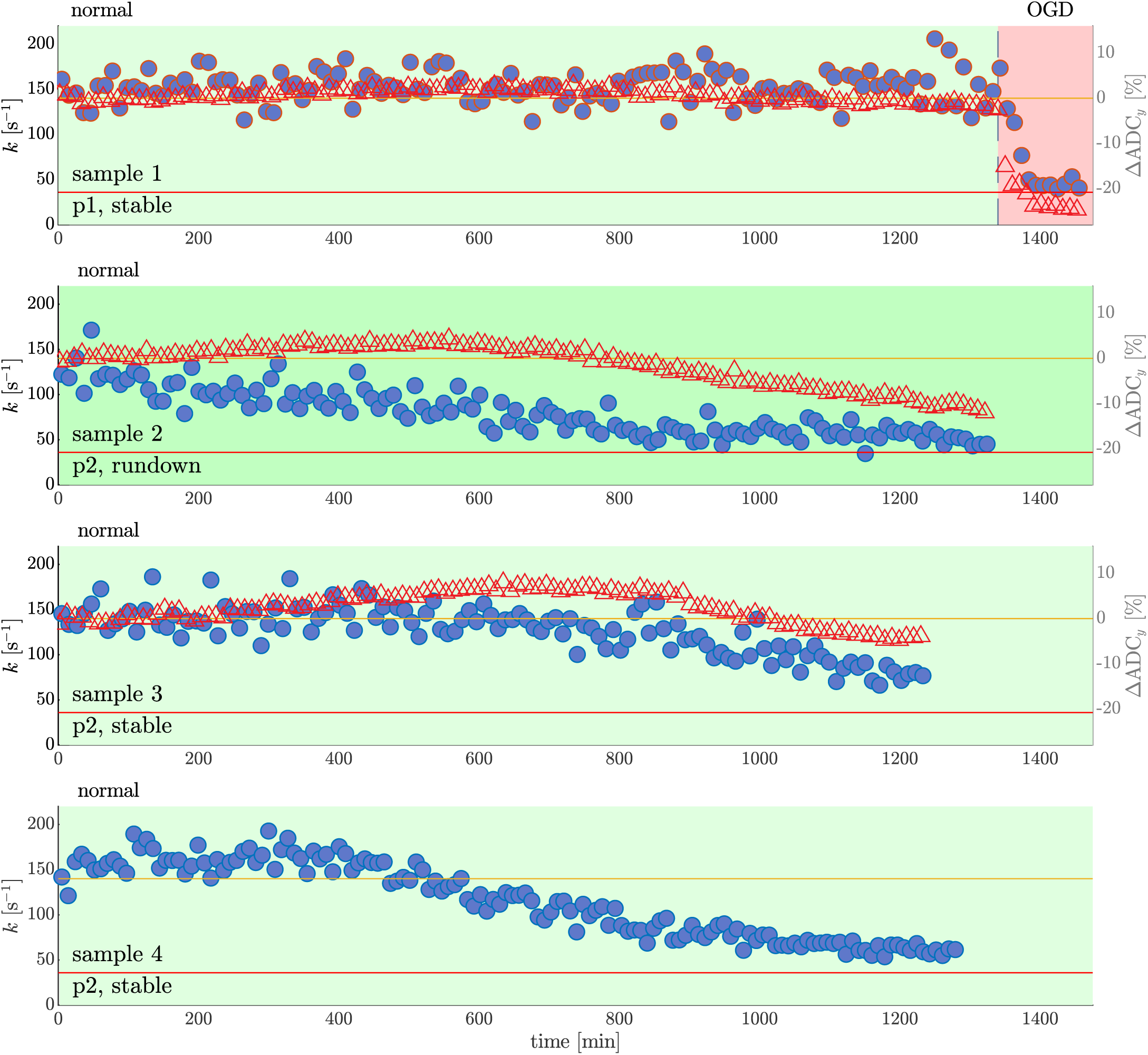
Stability tests. Exchange rates (solid blue circles, left axis) and normalized ADC_*y*_ changes (open red triangles, right axis) measured under normal conditions for extended periods for n=4 samples. For three of the samples, exchange rates remained stable for over six hours with values near the average from all samples under live conditions (*k* = 140 s^−1^, orange line). The backgrounds for the plots of these samples are shaded lighter. On sample 1, an OGD perturbation was performed at 22.3 hours. The exchange rate and ADC_*y*_ were stable over the entire period prior to OGD with mean ± SD 153 ±17 s^−1^ and 1.017 ±0.013µm^2^*/*ms respectively. After switching to OGD, the exchange rate decreased continuously for 40 minutes and then plateaued at 45 ±5 s^−1^, only slightly greater than the mean exchange rate of ouabain-treated samples (*k* = 36 s^−1^, red line). Exchange rates for the other two samples eventually ran down, after roughly 12 hours for sample 2 and 8 hours for sample 4, signifying an inevitable loss of viability. One sample (sample 3) started with four exchange rate values having a mean *<* 130 s^−1^. The background for the plot of this sample is shaded darker. If this sample was intended for an experiment, it would have been deemed unviable and discarded. Indeed, exchange rates for this sample continued to run down over the next 20 hours. Exchange rates during the last two hours of recording were 52 ±6 s^−1^. Diffusion was measured for samples 1–3. For sample 1, ADC_*y*_ values are stable until switching to OGD, at which point they decrease by 24%. For samples 2 and 3, ADC_*y*_ values increase prior to exchange rate rundown, and eventually decrease during exchange rate rundown. This non-monotonic behavior provides additional evidence that the ADC is not a direct measure of viability.

**Fig. S12.**
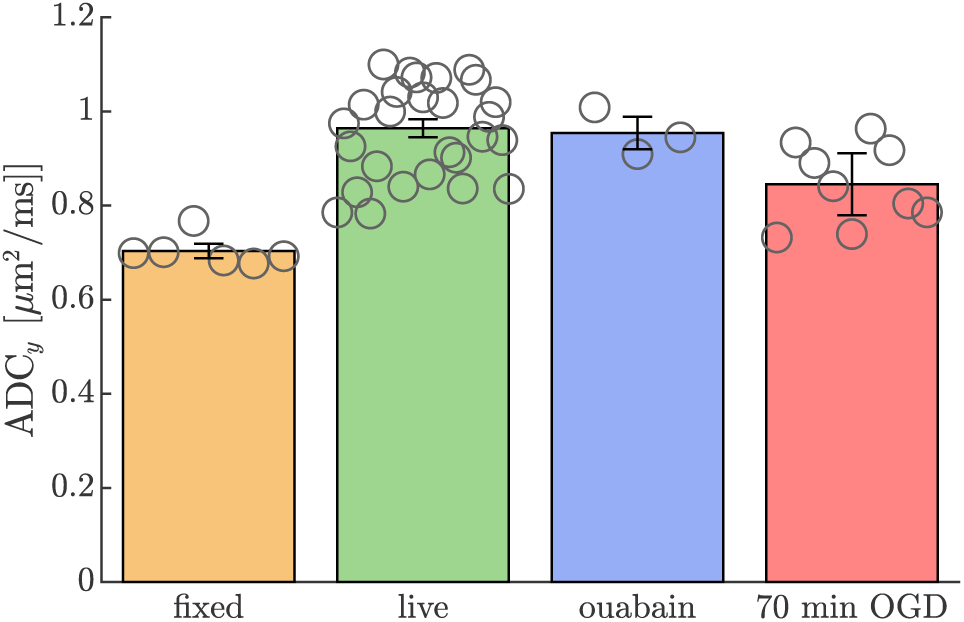
Apparent Diffusion Coefficients compared between fixed, live (untreated), ouabain-treated, and post-70 min OGD. Bar graphs present mean (bar height) 95% CI of the mean (whiskers), and average values from each sample (circles). ADC_*y*_ were compiled from the 1^st^ 25°C condition in Fig. 2, and from Figs. 3 and 4. The average values from each sample (open circles) are also shown. ADC_*y*_ of live spinal cords (mean ± SD 0.964 ± 0.097 *µ*m^2^*/*ms, *n* = 27) is significantly greater than fixed spinal cords (0.703 ± 0.031 *µ*m^2^*/*ms, *n* = 6) and spinal cords after 70 minutes of OGD (0.845 ± 0.085 *µ*m^2^*/*ms, n=9), (*p <* 0.001), but not significantly different from ouabain-treated spinal cords (0.954 ± 0.045 *µ*m^2^*/*ms, *n* = 3), (*p* = 0.75). Further, the ADC_*y*_ of ouabain-treated spinal cords is significantly greater than fixed spinal cords (*p <* 0.001)..

**Fig. S13.**
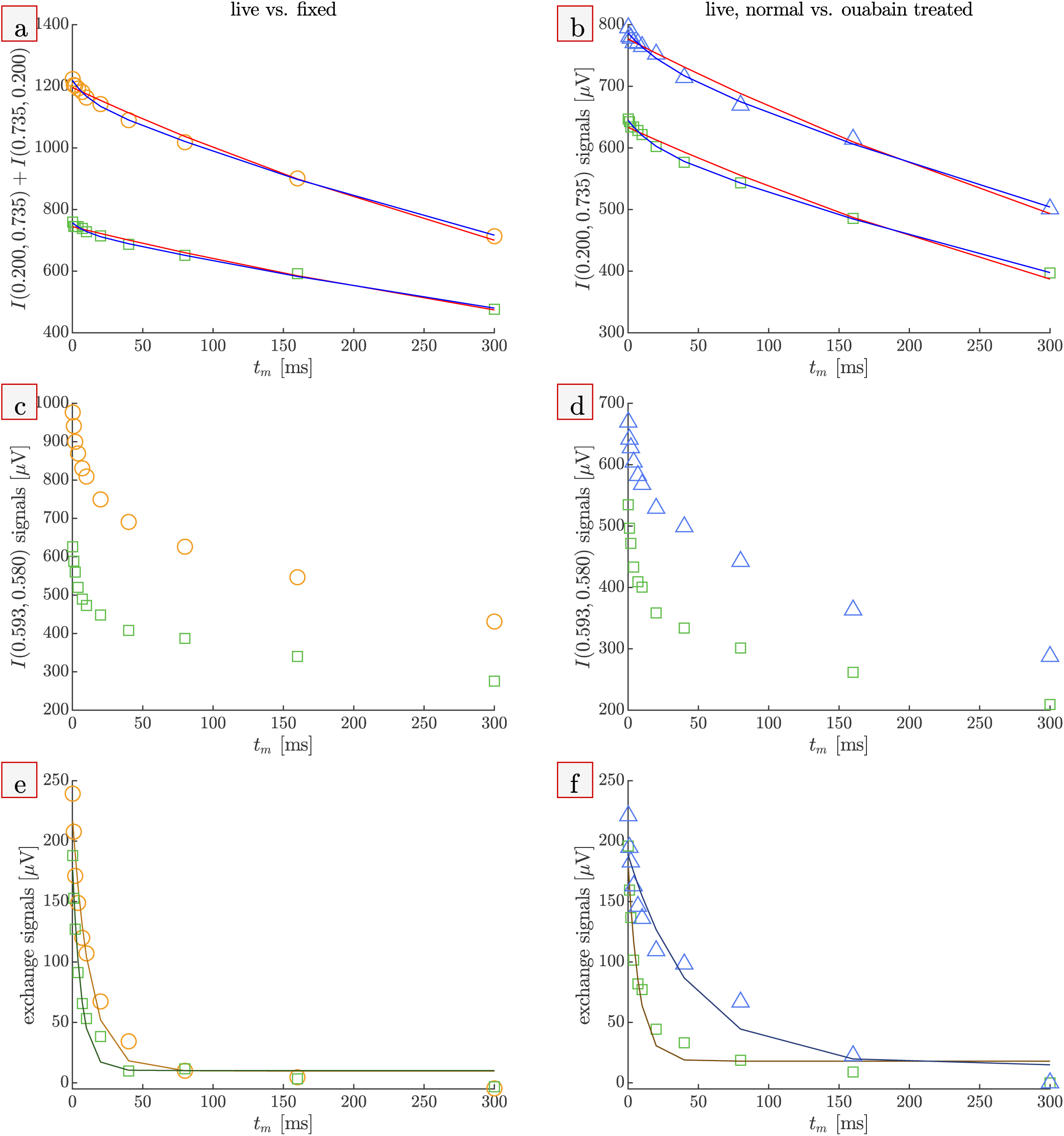
Representative DEXSY data and fits. Individual rapid exchange experiments acquired on (a, c, e) a live (green squares) and a fixed sample (orange circles) and (b, d, f) a live sample before (green square) and after (purple triangles) ouabain treatment. DEXSY signals were acquired as a function of mixing time with (*τ*_1_, *τ*_2_) combinations which make the signal (a and b) diffusion and *T*_1_ weighted but minimally exchange weighted and (c and d) diffusion, *T*_1_, and exchange weighted. a and b show representative single exponential fits (solid red lines) used to estimate diffusion weighted *R*_1_ and biexpoenential fits (solid blue lines) used to remove spin-lattice relaxation and isolate the “exchange signal”. e, f) Exchange signals, isolated by subtracting the biexponential fits shown in a and b from the signals in c and d. The signal at the final mixing time was subtracted from all signals so that the signals roughly decay to 0. Exchange rates were estimated by fitting a single exponential model with a baseline term to exchange signals. See Methods for details.

